# Profiling intra- and inter-individual differences in brain development across early adolescence

**DOI:** 10.1101/2022.12.19.521089

**Authors:** Katherine L. Bottenhorn, Carlos Cardenas-Iniguez, Kathryn L. Mills, Angela R. Laird, Megan M. Herting

## Abstract

As we move toward population-level developmental neuroscience, understanding intra- and inter-individual variability in brain maturation and sources of neurodevelopmental heterogeneity becomes paramount. Large-scale, longitudinal neuroimaging studies have uncovered group-level neurodevelopmental trajectories, and while recent work has begun to untangle intra- and inter-individual differences, they remain largely unclear. Here, we aim to quantify both intra- and inter-individual variability across facets of neurodevelopment across early adolescence (ages 8.92 to 13.83 years) in the Adolescent Brain Cognitive Development (ABCD) Study and examine inter-individual variability as a function of age, sex, and puberty. Our results provide novel insight into differences in annualized percent change in macrostructure, microstructure, and functional brain development from ages 9-13 years old. These findings reveal moderate age-related intra-individual change, but age-related differences in inter-individual variability only in a few measures of cortical macro- and microstructure development. Greater inter-individual variability in brain development were seen in mid-pubertal individuals, except for a few aspects of white matter development that were more variable between prepubertal individuals in some tracts. Although both sexes contributed to inter-individual differences in macrostructure and functional development in a few regions of the brain, we found limited support for hypotheses regarding greater male-than-female variability. This work highlights pockets of individual variability across facets of early adolescent brain development, while also highlighting regional differences in heterogeneity to facilitate future investigations in quantifying and probing nuances in normative development, and deviations therefrom.

## 1. Introduction

Recent years have seen great progress in charting child brain development, with much of the focus on developing normative models of structural change from a macroscale, or morphological, perspective (Aubert-Broche et al., 2013; Bethlehem et al., 2022; Mills et al., 2016; Tamnes et al., 2017; Wierenga et al., 2014). Convergent research has identified differential, curvilinear trajectories for morphometric aspects of macrostructural development, including cortical volume, thickness, and area, as well as subcortical and white matter volume from mid-childhood to early adulthood (Bethlehem et al., 2022; Herting & Sowell, 2017; Mills et al., 2016). On a more granular level, studies of brain microstructure reveal protracted white matter development into adolescence and through the mid-to late-twenties (Lebel & Deoni, 2018). Functional neuroimaging shows coherent large-scale brain networks in infants with topography similar to those of adults (Grayson & Fair, 2017), that are fine-tuned throughout childhood and adolescence (Cui et al., 2020). In this fine-tuning, functional topology of these networks’ changes, too, following trajectories of segregation and integration in the development of these large-scale brain networks (Fair et al., 2007; Marek et al., 2015). Therefore, there has been substantial progress in our understanding of neurodevelopment, although relatively less emphasis has been placed on understanding and modeling variability in these developmental processes (Becht & Mills, 2020), which is substantially different both within and between individuals (Ferschmann et al., 2022; Foulkes & Blakemore, 2018; Herting, Gautam, et al., 2018; Mills et al., 2021; Østby et al., 2009; Wierenga et al., 2014).

Dimensions of variability inherent to characterizing neurodevelopment include within-individual changes over time and between-individual differences in changes in brain maturation. Within-individual changes over time can vary across age and pubertal status, across regions of the brain, and across different facets of development (e.g., macroscale and microscale structural development, functional development) (Goddings et al., 2014; Herting, Johnson, et al., 2018; Tamnes et al., 2017). As such, our use of the term *intra-individual variability* is based on repeated assessments of brain MRIs that can be used to measure an individual’s developmental change. Such assessments have revealed normative within-individual variability underlying curvilinear development across wide-age ranges, such that rates of change within an individual differ as a function of increasing age and pubertal status (Aubert-Broche et al., 2013; Herting et al., 2017; Herting, Johnson, et al., 2018). This includes an inverted-U trajectory for gray matter volume with respect to age that peaks during childhood, a general increase in white matter volume across childhood and adolescence (Bethlehem et al., 2022; Paus et al., 2001), puberty-related nonlinearities in trajectories of both structural and functional development (Dai & Scherf, 2019; Goddings et al., 2014; Gracia-Tabuenca et al., 2021; Herting et al., 2015; Herting & Sowell, 2017). Regional variability in development includes differences in gray matter and white matter maturation (Aubert-Broche et al., 2013; Bethlehem et al., 2022; Mills et al., 2016), different developmental trajectories of cortical and subcortical gray matter (Bethlehem et al., 2022; Sowell et al., 2002), and differential timing of development in sensorimotor vs. association areas of the cortex, proceeding from phylogenetically older to newer regions (Gogtay et al., 2004). This intra-individual variability is foundational to studies of inter-individual variability and has implications for cognitive development (Battista et al., 2018; Blakemore & Choudhury, 2006), neurodevelopmental disorders (Grayson & Fair, 2017), and the emergence of psychopathology (Ferschmann et al., 2022; Paus et al., 2008).

Beyond changes within an individual, neurodevelopmental trajectories throughout late childhood and early adolescence vary between-individuals, as well. For example, inter-individual differences in brain outcomes are associated with physical and hormonal changes related to puberty (Dai & Scherf, 2019; Goddings et al., 2014; Herting & Sowell, 2017; Marceau et al., 2011; Vijayakumar et al., 2018), which also interact with sex- and age-related differences in neurodevelopment. Moreover, pubertal-, age-, and sex-related differences in development have implications for the neurobiology and emergence of psychopathology (Gogtay & Thompson, 2010; Graham et al., 2021; Paus et al., 2008; Shaw et al., 2010), which is more common during childhood and adolescence than other stages of life (Kessler et al., 2005; Solmi et al., 2022). Understanding associations between inter-individual differences in puberty and the brain is crucial to understanding development. However, much of the research into neurodevelopmental trajectories, and variability therein, focuses on brain macrostructure (e.g., cortical thickness, global and regional volume, white matter tract volume), while development of microstructure (e.g., cortical myelination, neurite density) and function (e.g., regional function and functional connectivity) has received comparatively little attention and is, thus, poorly characterized. Robust studies of individual differences in neuroimaging require substantially increased sample sizes (Button et al., 2013; Dick et al., 2021; Grady et al., 2021; Marek et al., 2022; Yarkoni, 2009), and are made possible by increasingly larger and more diverse samples with increased statistical power, capturing a broader range of individual variability. While developmental cognitive neuroimaging has made recent progress in profiling individual variability in brain development (Becht & Mills, 2020; Foulkes & Blakemore, 2018; Mills et al., 2021; Zhu & Qiu, 2022), this progress has been limited to small samples. There is still a long way to go in characterizing individual developmental trajectories and inter-individual differences in brain maturation over time.

While a few longitudinal studies have included and modeled subject-level estimates of brain development (Braams et al., 2015; Herting, Johnson, et al., 2018; Mills et al., 2021; Tamnes et al., 2017; van Duijvenvoorde et al., 2019; Wierenga et al., 2014), the focus remains largely on identifying mean trajectories. There is little work using variance to quantify individual differences during development, especially with the sample sizes needed to find robust effects. Studies either include individual-level change in presenting a group mean trajectory, as in the works mentioned above, or assess inter-individual variability in cross-sectional data. For example, Wierenga et al., who leveraged over 16,000 individuals’ MRI data from the ENIGMA Consortium to identify lifespan sex differences in variability across brain macrostructure measures (Wierenga et al., 2022). Individual variability in developmental change remains a gap in the field. Thus, additional research is necessary to characterize intra-individual neurodevelopmental change in a large sample as well as describe how age, sex, and pubertal maturation may contribute to inter-individual differences seen in changes of brain structure and function development. The Adolescent Brain Cognitive Development℠ Study (ABCD Study®) provides an unprecedented opportunity to study these important questions in the specific window of brain development that occurs during adolescence with tens of thousands of variables including multimodal neuroimaging measures and pubertal information acquired over multiple years (Jernigan & Brown, 2018). Such a large dataset has the potential to study intra- and inter-individual variability in developmental trajectories of brain macrostructure, microstructure, and function, and to elucidate inter-individual differences in brain development (Feldstein Ewing et al., 2018; Volkow et al., 2018). However, variability in these measures is likely not equally distributed. For example, there is greater variance in brain structure in boys than girls throughout childhood and adolescence (Wierenga et al., 2018, 2022) and greater variability across changes in brain structure during transitions into adolescence and adulthood than during childhood or mid-adolescence (Mills et al., 2021). For a more robust study of individual differences in brain development, researchers should consider not only the variability in developmental trajectories, but the homogeneity of this variability, i.e., heteroscedasticity. Assessing heteroscedasticity can reveal differences in brain change variability between levels of variables often included when studying brain development. Further, such insights can help identify pockets of variability for future study and inform sampling and experimental design for future research.

As such, this study presents a relatively novel approach to quantifying variability in neurodevelopmental trajectories with two overarching aims: to expand the current understanding of intra- and inter-individual variability in brain development across early adolescence from ages 9-13 years-old and to describe the distribution of this variability as a function of age, sex, and pubertal status. First, we examine within-person (i.e., intra-individual) variability in developmental trajectories in brain macrostructure, microstructure, and function by calculating annual percent change across the brain in each individual. Then, we investigate between-person (i.e., inter-individual variability in developmental trajectories by computing the variance in brain changes across the whole sample. Finally, we assess the distribution of between-individual variability in neurodevelopment by assessing heteroscedasticity in brain changes across age, sex, and pubertal stages. In characterizing within-individual, or intra-individual, developmental change we have chosen to calculate annualized percent change (APΔ) for any given measure of brain structure or function. This approach has several advantages. First, computing change within each individual participant estimates the slope of an individual’s developmental trajectory between two time points, mitigating the effect of differences in average values between individuals (Mills et al., 2021). Second, annualized percent change removes the effect of measurement scale or brain region size, allowing comparisons across neuroimaging measures (e.g., microstructure, functional connectivity). Third, normative studies of child brain development indicate that multiple, overlapping neurobiological processes occur between ages 9-13 years, including myelination, synaptic pruning, synaptogenesis, and apoptosis (Tau & Peterson, 2010) and associated changes follow different trajectories across the brain. By modeling annualized percent change in a variety of neuroimaging measures, we can capture intra-individual differences in these trajectories across brain regions, tissue types, and facets of development,providing broadly applicable information about the inherent variability in development occurring during this important period of early adolescence. Finally, annualized percent change mitigates the extent to which differences in elapsed time between assessments across individuals confound estimates during this period of such rapid and varied neurodevelopment. Using annualized percent change, we investigated heterogeneity in intra-individual differences in macrostructural, microstructural, and functional brain changes from ages 9 to 13 years, and complement the well-developed literature on normative macrostructural development with an innovative approach to studying variability with respect to age, sex, and puberty. We chose to include a broad swath of neuroimaging measures for a comprehensive look at inter-individual differences in neurodevelopment, to focus on puberty for its ubiquitous impact on ABCD Study participants and its interest to the broad research community using ABCD Study data. In doing so, we hope to shed new light on neurodevelopmental trajectories during the transition from late childhood to early adolescence.

## 2. Methods

### 2.1. Participants

Longitudinal data were collected as a part of the ongoing Adolescent Brain and Cognitive Development (ABCD) Study, and included in the annual 4.0 data release (http://dx.doi.org/10.15154/1523041). The ABCD Study enrolled 11,880 children 9 to 10 years of age (mean age = 9.49; 48% female) in a 10-year longitudinal study. Participants were recruited at 21 study sites across the United States from elementary schools (private, public, and charter schools) in a sampling design that aimed to represent the nationwide sociodemographic diversity (Garavan et al., 2018). All experimental and consent procedures were approved by the institutional review board and human research protections programs at the University of California San Diego. ABCD Study inclusion criteria included age at enrollment (9.0 to 10.99 years); English fluency; lack of MRI contraindications; no history of traumatic brain injury or major neurological disorder; absence of any non-correctable sensory and/or motor impairments that would preclude the youth’s participation in study procedures; current intoxication at appointment; diagnosis of any DSM-I psychotic, autism spectrum, or substance use disorder; an intellectual disability reported by their caregiver; premature birth, very low birth weight, or perinatal complications; and caregiver knowledge at baseline of an impending move to an area beyond reasonable traveling distance to an ABCD Study site. Each participant provided written assent to participate in the study and their legal guardian provided written agreement to participate. For more information, see Garavan et al. (2018) and Volkow et al. (Volkow et al., 2018). Here, we use a subset of data from the ABCD Study, including magnetic resonance imaging (MRI), in addition to variables regarding participants’ age at time of study enrollment, sex at birth and pubertal development. Data include assessments from two time points: baseline enrollment (i.e., at ages ∼9-10 years) and year 2 follow-up (i.e., at ages ∼11-13 years). Participants’ sex was taken from the baseline time point, while pubertal stage was considered at baseline, year 1 follow-up, and year 2 follow-up.

Participants were excluded from these analyses if they did not have imaging data collected at the 2-year follow-up visit. From this subset, data were further excluded based on image quality. Structural and diffusion-weighted data were included if a participant’s T1- and diffusion-weighted images, respectively, met the ABCD-recommended criteria for inclusion. For T1-weighted images, quality assessments were based on motion, intensity inhomogeneity, white matter and pial surface estimation by Freesurfer, and susceptibility artifacts and exclusion was recommended if an image exhibited severe artifact in any of those categories. For diffusion-weighted images, quality assessments were based on residual B_0_ distortion after processing, coregistration to the participant’s T1, image quality, and segmentation quality, with exclusion recommended if an image exhibited severe artifact in any of those categories. For resting-state scans, quality assessments were based on the number of frames remaining after high-motion (i.e., framewise displacement > 0.3 mm) frames were censored and periods with fewer than 5 contiguous frames were excluded, coregistration to T1 success, B_0_ distortion maps, Freesurfer tissue type segmentation quality, and the number of usable time points per run (after censoring and exclusion, runs with fewer than 100 usable time points were excluded). We further excluded any resting-state fMRI data from participants with fewer than 750 usable (i.e., low-motion) frames, or 10 low-motion minutes, across all runs. This decision was based on estimations of scan lengths necessary for reliable resting-state functional connectivity estimates (Birn et al., 2013; Noble et al., 2017). The final sample characteristics for the current study are described in Table 1. Because each structural, diffusion-weighted, and resting-state functional data are included based on their image quality, there are different numbers of subjects included for analyses with different imaging modalities. Those values are in Supplemental Table 2.

**Table 1:** Sample Demographics

|  | Full ABCD Study Sample |  | MRI 2-Year Follow-Up |  |
| --- | --- | --- | --- | --- |
|  | N | % of sample | N | % of sample |
| Total N | 11801 |  | 7457 |  |
| **Age at baseline (months)** | 118.97±7.49 |  | 118.74±7.44 |  |
| **Sex (F)** | 5636 | (48%) | 3437 | (46%) |
| ***Race & Ethnicity*** |  |  |  |  |
| Other | 1493 | (13%) | 918 | (12%) |
| Hispanic | 2402 | (20%) | 1450 | (19%) |
| Black | 1755 | (15%) | 981 | (13%) |
| White | 6149 | (52%) | 4108 | (55%) |
| ***Household Income*** |  |  |  |  |
| < \$75,000 | 3194 | (27%) | 1922 | (26%) |
| \$75,000 to \$100,000 | 3056 | (26%) | 2082 | (28%) |
| > \$100,000 | 4543 | (38%) | 2876 | (39%) |
| ***Caregiver Education*** |  |  |  |  |
| Up to high school diploma, GED | 2022 | (17%) | 1147 | (15%) |
| Some college, associate degree | 3464 | (29%) | 2233 | (30%) |
| Bachelor's degree | 3317 | (28%) | 2204 | (30%) |
| Graduate degree | 2981 | (25%) | 1863 | (25%) |
| **Caregiver Marital Status** |  |  |  |  |
| Married | 7951 | (67%) | 5192 | (70%) |
| Widowed | 96 | (<1%) | 60 | (<1%) |
| Divorced | 1077 | (9%) | 659 | (9%) |
| Separated | 461 | (4%) | 262 | (4%) |
| Never Married | 1444 | (12%) | 850 | (11%) |
| **MRI Scanner Manufacturer** |  |  |  |  |
| Siemens | 7303 | 62% | 4539 | 61% |
| GE Medical Systems | 2977 | 25% | 2013 | 27% |
| Philips Medical Systems | 1521 | 13% | 905 | 12% |

*Note.* Differences in baseline demographic composition of the complete sample and the sample including individuals with high-quality MRI data collected at a 2-year follow-up appointment were assessed using Mann-Whitney U tests. Bold categories indicate significant differences at $\alpha < 0.05$ ; bold and italicized categories, at $\alpha < 0.01$ . Within-category percentages that add to <100% are due to missing data and participants who declined to answer, i.e., in the case of household income. *MRI 2-Year Follow-Up* indicates the sample used in analyses presented here. The "Other" Race/Ethnicity category includes participants whose caregiver identified them as American Indian/Native American, Alaska Native, Native Hawaiian, Guamanian, Samoan, Other Pacific Islander, Asian Indian, Chinese, Filipino, Japanese, Korean, Vietnamese, Other Asian, Other Race, or as belonging to more than one race.

### 2.2. Developmental variables and imaging measures

For clarity, “variables” will be used to refer to non-brain assessments of development and demographics (i.e., age, sex, pubertal status) (i.e., age, sex, pubertal status) and “measures” will be used to refer to brain imaging outcomes outcomes throughout.

#### 2.2.1. Puberty

Pubertal status was calculated from the Pubertal Development Scale (Petersen et al., 1988), completed by the caregiver about the participant (Barch et al., 2018; Herting et al., 2021). The PDS consists of a total of 5 distinct questions for males and females, based on sex assigned at birth. For males, the five questions assess height growth, body hair, skin changes, vocal changes, and facial hair. For females, the five questions consist of assessing changes in height growth, body hair, skin, breast development, and menarche. For each of the 5 questions, parents/caregivers and youth were asked to separately rate their development on a 4-point scale (1 = has not begun yet, 2 = barely begun, 3 = definitely begun, 4 = seems complete), except for the menarche question for females, which consisted of a yes/no answer choice. Each question also consisted of an “I don’t know” answer. A pubertal category score was derived for male participants by summing the body hair growth, voice change, and facial hair items and categorizing them as follows: prepubertal = 3; early pubertal = 4 or 5 (no 3-point responses); mid-pubertal = 6-8 (no 4-point responses); late pubertal = 9-11; postpubertal = 12. The puberty category score was derived for female participants by summing the body hair growth, breast development, and menarche and categorizing them as follows: prepubertal = 3; early pubertal = 3 and no menarche; midpubertal = 4 and no menarche; late pubertal<=7 and menarche; postpubertal= 8 and menarche. These category scores roughly correspond to Tanner stages and fall on a scale from 1 (prepubertal) to 5 (post pubertal) (Cheng et al., 2021; Herting et al., 2021).

The distribution and characteristics of pubertal assessments in the ABCD Study at baseline are profiled by Herting et al. (2021). In the current analytic sample, a greater proportion of male participants are “prepubertal” at baseline data collection (i.e., ages 9-10 years) than are female participants (Supplemental Figure 1).

#### 2.2.2. Neuroimaging Data

The ABCD Study’s imaging protocol includes structural, diffusion, and both task-based and resting-state functional MRI collected every two years, as described by Casey et al. (2018). The ABCD consortium has reported image processing and analysis methods in detail (Hagler et al., 2019). Important for multi-site studies, ABCD MRI methods and assessments have been optimized and harmonized across the 21 sites for 3 Tesla scanners (Siemens Prisma, General Electric 750, Philips) (Casey et al., 2018; Hagler et al., 2019). For more details, see Supplemental Methods.

Here, we incorporated a range of imaging measures across these MRI modalities for a more comprehensive assessment of brain development. Broadly, these measures assess brain morphometry (i.e., macrostructure), microstructure, and function, and are described below in Table 2. Imaging measures were included for one of two reasons: either because they are widely researched in the developmental neuroscience literature (e.g., cortical thickness, area, fractional anisotropy) or they are relatively novel in developmental neuroscience and offer additional insight to the neurobiology of development not addressed by the other included measures (e.g., intracellular diffusion, BOLD variability).

**Table 2:** Neuroimaging Measures of Interest

| Modality | Measures | Description |
| --- | --- | --- |
| Macrostructure |  |  |
| sMRI | Cortical volume | Estimated volume of each cortical region of interest (ROI) (Durham et al., 2021) |
|  | Cortical area | Surface area of each cortical region of interest (ROI; (Dale et al., 1999)) |
|  | Cortical thickness | Thickness of the cortical ribbon (i.e., from the pial surface to gray / white matter boundary) of each cortical ROI (Fischl & Dale, 2000) |
|  | Subcortical volume | Volume of each subcortical ROI (Fischl et al., 2002) |
| dMRI | White matter tract volume | Volume of each white matter tract (Hagler et al., 2009) |
| Microstructure |  |  |
| sMRI | Gray matter / white matter contrast | Contrast of T1-weighted image intensity across gray-white matter boundary in cortical ROIs, indicative of neural tissue properties (Salat et al., 2009) |
| dMRI – RSI | Isotropic intracellular diffusion* | Magnitude of spherical intracellular diffusion, as if in a cell body; within each white matter tract, cortical and subcortical ROI (White et al., 2012) |
|  | Directional intracellular diffusion* | Magnitude of directional intracellular diffusion, as if in an axon or dendrite; within each white matter tract, cortical and subcortical ROI (White et al., 2012) |
| dMRI – DTI | Fractional anisotropy | Degree to which diffusion follows a single direction within each white matter tract, a proxy for cellular microstructure (Alexander et al., 2007; Clark et al., 2011) |
|  | Mean diffusivity | Magnitude of diffusion within each white matter tract, estimates the restriction of water movement in tissue (Alexander et al., 2007; Clark et al., 2011) |
|  | Transverse diffusivity | Average of the magnitude of diffusion in secondary and tertiary diffusion directions (i.e., second and third eigenvalues; Alexander et al., 2007) |
|  | Longitudinal diffusivity | Magnitude of diffusion in primary diffusion direction (i.e., first eigenvalue; Alexander et al., 2007) |
| Function |  |  |
| rs-fMRI | BOLD temporal variance | Variance in the average blood-oxygen-level-dependent (BOLD) signal across each cortical and subcortical ROI resting-state fMRI scans, reflecting the magnitude of low-frequency BOLD oscillations (Wang et al., 2021) |
|  | Between-network functional connectivity | Pairwise correlations between BOLD signals from each large-scale brain network (Gordon et al., 2016) |
|  | Within-network functional connectivity | Average BOLD signal correlation of regions within each large-scale brain network (Gordon et al., 2016) |
|  | Network-subcortical functional connectivity | Pairwise correlations between BOLD signals from each large-scale brain network (Gordon et al., 2016) and each subcortical ROI |

*Note.* Cortical regions of interest are defined using automated FreeSurfer parcellations (Desikan et al., 2006), including 68 cortical regions. Subcortical regions of interest are defined using automated FreeSurfer parcellations based on the Harvard-Oxford probabilistic atlas (Fischl, 2012; Fischl et al., 2002), from which 22 subcortical structures are included here. White matter tracts are defined using AtlasTrack (Hagler et al 2009), from which 35 tracts are included here. Cortical networks are functionally defined (Gordon et al., 2016), comprising 13 canonical networks and a grouping of extra-network regions for a total of 14 parcels. \*Isotropic intracellular diffusion corresponds to restricted normalized isotropic diffusion in the ABCD Study data dictionaries; directional intracellular diffusion, to restricted normalized directional diffusion. This is reflected in their abbreviation (RNI, RND) in figures throughout the manuscript.

### 2.3. Analyses

An analysis plan was registered for this project with the <u>Open Science Framework (OSF</u>; https://doi.org/10.17605/OSF.IO/ZCRW8). Prior to data analysis, missingness was assessed across imaging measures and demographic variables and for the purposes of assessing intra- and inter-individual variability only complete cases were used. As the number of participants with high-quality MRI data differed based on the scan (i.e., T1- and diffusion-weighted, resting-state functional), assessments of imaging measures comprised slightly different numbers of participants. A total of 7547 participants had data from the 2-year follow-up visit, 7115 (95%) of these had high-quality T1-weighted, structural data from both baseline and 2-year follow-up visits (i.e., assessing cortical thickness, volume, area, and gray/white matter contrast), 6248 (84%) of these had high-quality diffusion-weighted data (i.e., assessing fractional anisotropy; mean, longitudinal, and transverse diffusivity; directional and isotropic intracellular diffusion; white matter tract volume), and 4119 (55%) of these had high-quality resting-state functional data (i.e., assessing BOLD variance, within-/between-network connectivity, and subcortical-network connectivity). Further details can be found in Supplementary Table 2.

#### 2.3.1. Intra-individual change

Intra-individual changes in brain measures were modeled as annualized percent change, calculated by dividing the percent change in each measure by the elapsed time between observations, per the following formula (Mills et al., 2021):

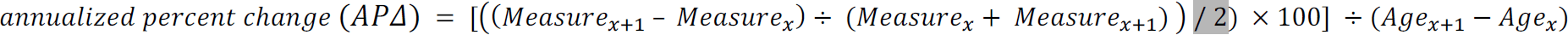

Given prior findings by our group that an individual’s developmental trajectory is related to their starting point (Mills et al., 2021), we further computed partial correlation coefficients between the initial values of each brain feature at study enrollment, ages at study enrollment (i.e., between 9 and 10 years), and APΔ for each brain measure. Specifically, we performed sets of partial correlation analyses to disentangle the unique, group-level associations between 1) age at study enrollment and APΔ and 2) initial values of each brain feature (at the first wave of data collection) and APΔ.

#### 2.3.2. Inter-individual differences in change

To assess inter-individual differences in brain development, distributions of intra-individual change were, then, compared across levels of the following developmental variables using the Fligner-Killeen test for homogeneity of variances (Fligner & Killeen, 1976): participants’ sex assigned at birth; pubertal status (i.e., pubertal category score from the Pubertal Development Scale); and age at study enrollment (across 4 equal-sized bins: 9.0 – 9.39 years old, 9.4 – 9.99 years old, 10.0 – 10.49 years old, 10.5 – 10.99 years old). Heteroscedasticity (i.e., inhomogeneity of variance) across age bins assesses if inter-individual differences vary based on an individual’s age, which is in contrast to the correlations and partial correlations described in section 2.3.1 that examines if the magnitude and direction of intra-individual change relates to an individual’s age, across the sample.

In our sample, distributions of participants in “prepubertal”, “early puberty”, and “midpubertal” stages are approximately equal (albeit with different proportions of male and female participants within each stage), with <100 participants considered “late pubertal” (Supplemental Figure 1). Thus, heteroscedasticity was assessed between participants in “prepubertal”, “early puberty”, and “midpubertal” stages.

Significance was assessed at *α_adjusted_* < 0.05, corrected for the number of effective comparisons across APΔ per measure, per region (Li & Ji, 2005; Šidák, 1967). This approach used all 1144 APΔ values (each region’s value per measure) and uses the eigenvalues of a matrix of pairwise correlations between APΔ measures to determine the number of effective comparisons to correct for, thus accounting for dependence between measures. The Li & Ji adjustment to the Šidák correction to control familywise error rate revealed 534.5 “effective” comparisons across all 1144 APΔ measures, adjusting *α* < 0.05 to *α* < 0.000096. This threshold was used as the significance threshold for all reported *p*-values.

Then, post hoc rank-sum tests were performed for brain change across significantly heteroscedastic measures, per developmental variable (i.e. age bins, sex, pubertal stages) and values of APΔ were plotted across age bins, sex at birth, and pubertal stages to visualize inter-individual differences. Finally, patterns were assessed across measures of brain change according to (a) which biological concepts (i.e., macrostructure, microstructure, and function) exhibited the greatest difference in variance across developmental variables, and (b) which brain regions exhibited the greatest differences in variance.

All code used to perform these analyses and generate the associated figures was written in Python 3.7 and is available at github.com/62442katieb/deltaABCD_variability. A mapping between those scripts and the relevant analyses, figures, and tables is provided in Supplemental Table 0.

#### 2.3.3. Additional exploratory analyses

To more fully explore the role of puberty in inter-individual variability, we assessed pubertal stage across the 2-year span between imaging data acquisitions. In this span, participants’ caregivers completed the PDS annually (i.e., at ages ∼9-10, ∼10-11, and ∼11-12 years). These distributions of pubertal stage were examined by age and sex (Supplemental Figure 1). Unlike sex at birth, which is time invariant, pubertal stage varies by sex at birth, and changes over time. Thus, we also examined changes in pubertal stage between waves of data collection and compared variance in each imaging variable across pubertal stages at each wave of data collection, and across changes in pubertal stage between waves, using the Fligner-Killeen test for homogeneity of variances. These analyses were repeated for male and female participants, separately, due to sex differences in pubertal stages in the sample.

## 3. Results

Our results reveal differences in the distributions of *intra*-individual change across brain regions, white matter tracts, and networks based on both features of brain development (i.e., macrostructure, microstructure, function) and tissue types (i.e., gray and white matter). Further, greater *inter*-individual variability is seen in changing function and functional connectivity than in either macro- or microstructural changes. Further, assessments of heteroscedasticity indicate that inter-individual variability in neurodevelopmental change is unequally distributed across participants’ age at enrollment (i.e., 9-10 years), sex at birth, or pubertal status. Below we report greater details regarding patterns of intra-individual variability in annual percent change (APΔ) across the brain, as well as inter-individual differences in these changes observed between the ages of 9-13 years.

### 3.1. Intra-individual changes in brain development

Intra-individual differences as measured by APΔ are presented for measures of macrostructure, microstructure, and brain function in Figure 1 and Table 3. Consistent with extant literature on this age range, cortical thickness and gray matter volume decreased annually across brain regions, while white matter volume increased. Likewise, measures of white matter organization (i.e., fractional anisotropy) largely increased, while magnitudes of diffusion (i.e., longitudinal, transverse, and mean diffusivity) decreased. However, isotropic intracellular diffusion increased in both gray and white matter regions, while directional intracellular diffusion increased in white matter tracts but decreased in gray matter regions. Moreover, both isotropic and directional intracellular diffusion exhibited greater variability in gray matter than in white matter. Finally, while macro- and microstructural measures largely either decreased or increased across the brain (with the exception of cortical area), measures of brain function exhibited both positive and negative APΔ. Finally, standard deviations of functional measures were an order of magnitude larger, on average, than those of macrostructure and microstructure measures.

**Figure 1.**
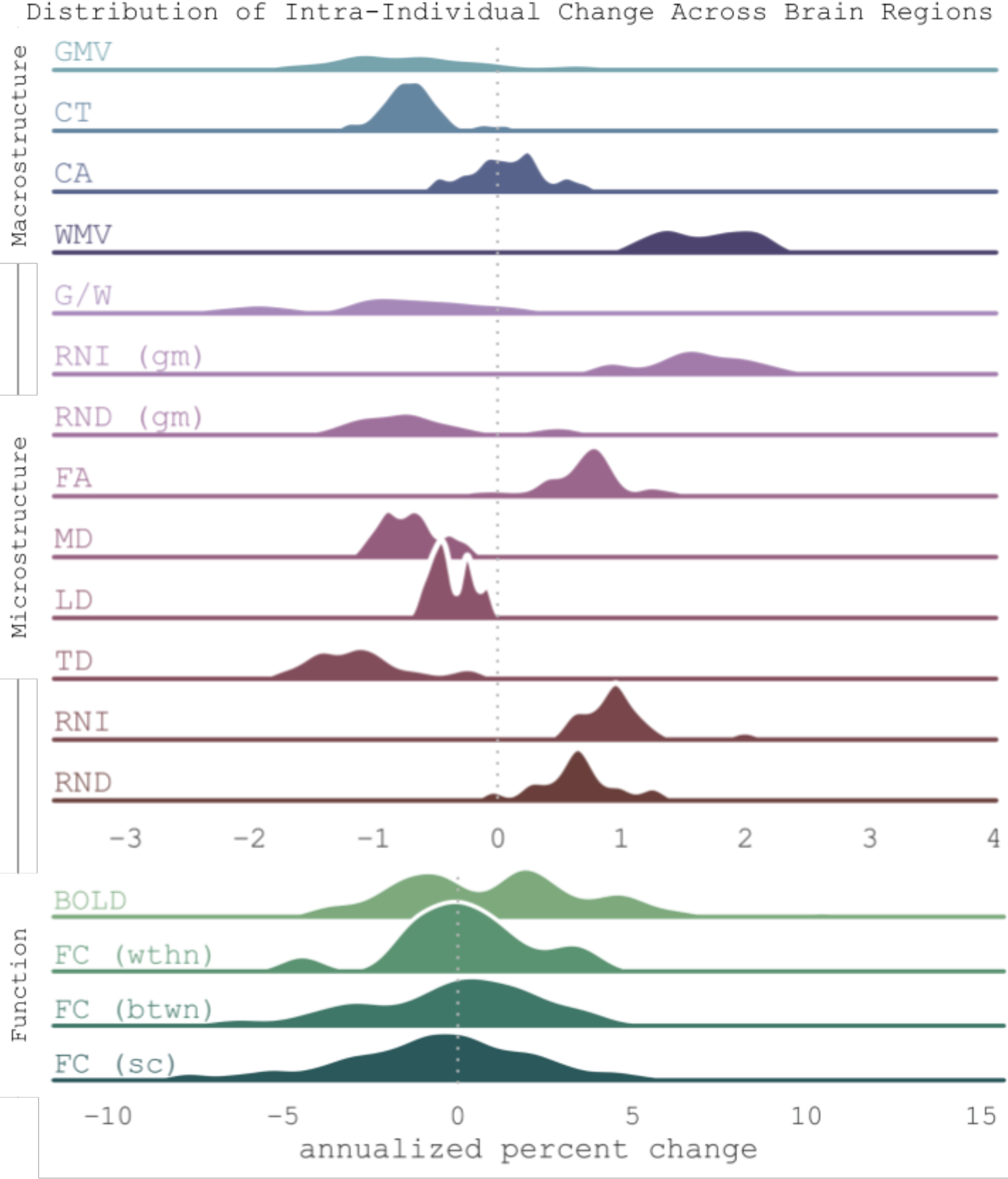
Distributions of annualized percent change in brain measures summarized across all participants and brain regions, separated by neuroimaging measure and color-coded by biological construct the measure represents. In order, measures of macrostructure include gray matter volume (GMV), cortical thickness, cortical area, and white matter volume (WMV); measures of microstructure include gray-to-white matter contrast (GM/WM contrast), mean diffusivity (WM MD), fractional anisotropy (WM FA), isotropic intracellular diffusion of white matter and both cortical and subcortical gray matter (WM RNI, GM RNI), directional intracellular diffusion of white matter and both cortical and subcortical gray matter (WM RND, GM RND); and measures of brain function include blood-oxygen-level-dependent (BOLD) temporal variance (BOLD Variance), cortical within-network functional connectivity (Within FC), cortical between-network functional connectivity (Between FC), subcortical-network connectivity (Subcortical FC). Dotted line marks zero point, indicating no change per year, whereas positive values reflect increases and negative values represent decreases in percent change per year.

**Table 3:** Annualized percent change in neurodevelopment summarized across brain regions per neuroimaging measure

| | MRI Measure | $\bar{x}_{AP\Delta}$ | $\sigma_{AP\Delta}$ | [Q<sub>1</sub>, Q<sub>3</sub>] |
| --- | --- | --- | --- | --- |
| Macrostructure | GM volume | -0.49 | 2.56 | (-1.12, -0.06) |
|  | Cortical area | 0.08 | 2.34 | (-0.12, 0.28) |
|  | Cortical thickness | -0.68 | 1.71 | (-0.85, -0.55) |
|  | WM volume | 1.72 | 2.47 | (1.37, 2.05) |
|  | Gray-to-white matter contrast | -0.93 | 3.71 | (-1.57, -0.38) |
| Microstructure | Isotropic intracellular diffusion (GM) | 1.54 | 2.89 | (1.31, 1.95) |
|  | Directional intracellular diffusion (GM) | -0.65 | 4.45 | (-1.06, -0.35) |
|  | Fractional anisotropy (WM) | 0.69 | 2.04 | (0.52, 0.87) |
|  | Mean diffusivity (WM) | -0.7 | 1.84 | (-0.89, -0.57) |
|  | Transverse diffusivity (WM) | -1.09 | 1.92 | (-1.4, -0.94) |
|  | Longitudinal diffusivity (WM) | -0.37 | 1.79 | (-0.5, -0.25) |
|  | Isotropic intracellular diffusion (WM) | 0.98 | 1.55 | (0.79, 1.07) |
|  | Directional intracellular diffusion (WM) | 0.64 | 1.61 | (0.49, 0.79) |
| Function | BOLD variance | 1.18 | 22.73 | (-0.99, 2.69) |
|  | Within-network FC | 0.18 | 12.25 | (-0.94, 1.05) |
|  | Between-network FC | 0.05 | 41.66 | (-1.7, 1.84) |
|  | Subcortical-network FC | -0.71 | 47.28 | (-2.4, 1.23) |

*Note.* Values represent average mean ( $\bar{x}$ ) and standard deviation ( $\sigma$ ), and 1st and 3rd quartiles (Q<sub>1</sub>, Q<sub>3</sub>) of annualized percent change ( $AP\Delta$ ) across brain regions. Abbreviations: GM = gray matter; WM = white matter; FC = functional connectivity; BOLD = blood-oxygen-level-dependent.

#### 3.1.1. Neuroanatomy of intra-individual changes

Within individuals, trends in annualized percent change varied neuroanatomically (Figure 2 and supplemental results).

**Figure 2.**
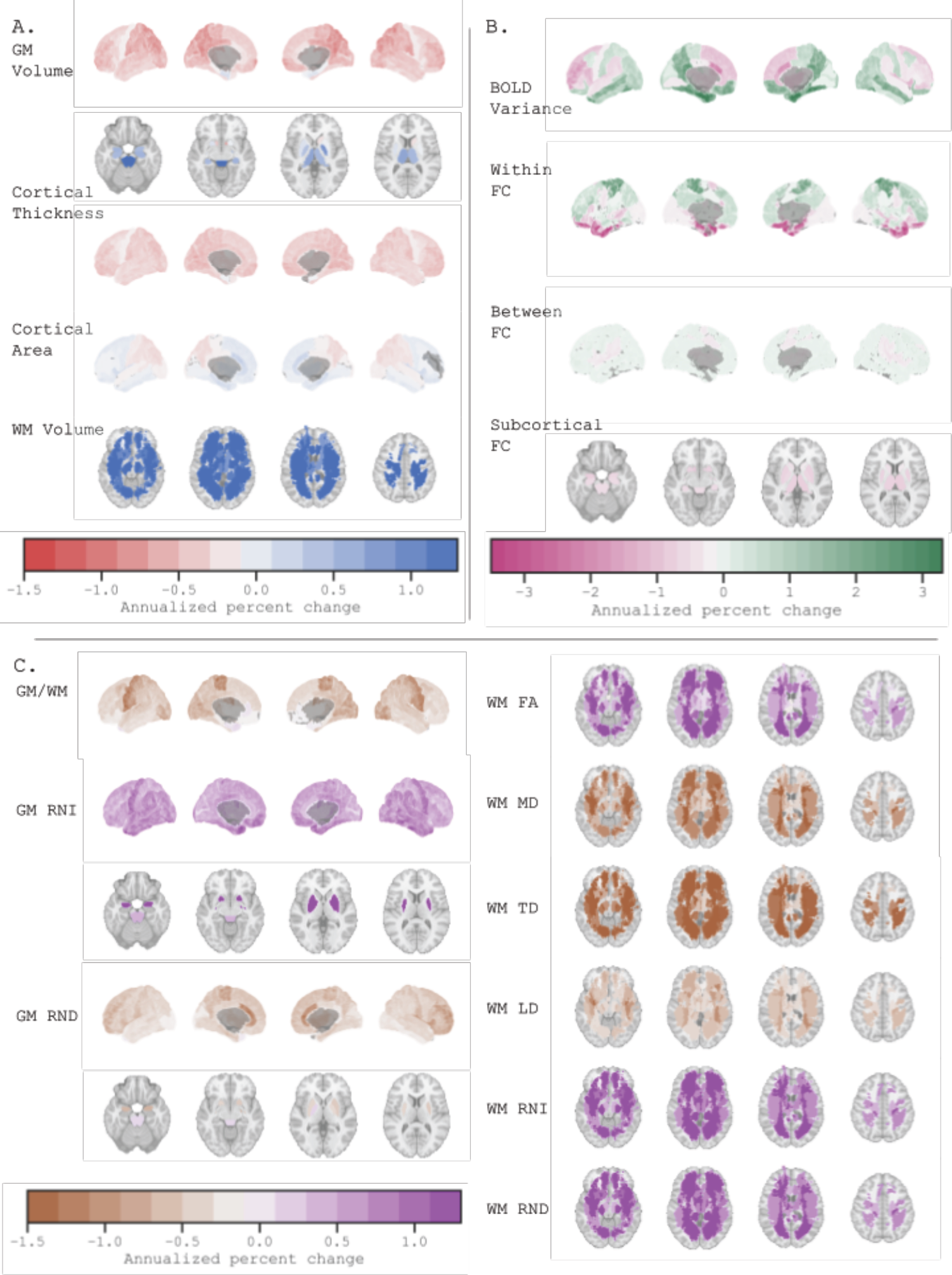
Average annualized percent change in brain measures across participants, separated by measure and color-coded by biological construct the measure represents. A) Macrostructure annualized percent change, including gray matter volume (GM Volume), cortical thickness, cortical area, and white matter volume (WM Volume). B) Brain function annualized percent change, including BOLD temporal variance (BOLD Variance), cortical within-network functional connectivity (Within FC), cortical between-network functional connectivity (Between FC), subcortical-network connectivity (Subcortical FC). Volumetric files of APΔ across the brain, per measure, are available on the Open Science Foundation (https://osf.io/a9kcv/). C) Gray matter (GM) and white matter (WM) microstructure change annualized percent change, including gray-to-white matter contrast (GM/WM), both cortical and subcortical gray matter isotropic intracellular diffusion (GM RNI), both cortical and subcortical gray matter directional intracellular diffusion (GM RND), white matter fractional anisotropy (WM FA), white matter mean diffusivity (WM MD), white matter transverse diffusivity (WM TD), white matter longitudinal diffusivity (WM LD), white matter isotropic intracellular diffusion (WM RNI), white matter directional intracellular diffusion (WM RND). Abbreviations: L = left; R = right; BOLD = blood-oxygen-level-dependent. Subcortical slices: z = [-20, -10, 0, 10]; white matter slices = [-10, 3, 18, 40].

Across *macrostructural* measures (Figure 2A), the greatest decreases were seen in posterior regions’ cortical volume and thickness, while cortical area exhibited posterior decreases and anterior increases. White matter volume increases were relatively uniform across tracts, while subcortical volume increases were greatest in the brainstem.

Across *microstructural* measures (Figure 2C), the greatest decreases were seen in pre- and postcentral gyrus gray-to-white matter contrast, anterior cingulate directional intracellular diffusion, and diffusivity of bilateral fornix, striatum, anterior thalamic radiations, and inferior fronto-occipital fasciculus. Greatest gray matter microstructure increases were seen in isotropic intracellular diffusion of precentral, temporal, and subcortical (i.e., pallidum, thalamus, nucleus accumbens, amygdala) regions. The largest white matter microstructure increases were seen in bilateral parahippocampal cingulum and uncinate fasciculus. White matter fractional anisotropy increased most across the cingulum, superior longitudinal and inferior fronto-occipital fasciculus.

Across *functional* changes (Figure 2B), the greatest increases were seen in BOLD variance of bilateral entorhinal cortices, fusiform gyri, and precuneus, as well as within-network connectivity of the sensorimotor hand and cinguloparietal networks. Between-network connectivity saw a mix of increases and decreases, depending on the connection, with no clear increases or decreases across most of any one network’s connections. The greatest functional decreases were seen in BOLD variance of the caudal anterior cingulate, dorsolateral frontal, and cerebellar cortices, as well as cortical connectivity of the putamen and thalamus, and connectivity of extra-network regions.

#### 3.1.2. Group-level associations between changes, initial values of brain features, and age at study enrollment

Partial correlations between APΔ and each (i) initial values of brain features at study enrollment (i.e., first wave of data collection) and (ii) participant age at study enrollment allowed us to assess unique start-change and age-change associations across participants, respectively. Overall, start-change correlations were an order of magnitude larger than age-change correlations

Associations between initial values were overwhelmingly significant and largely negative across measures and brain regions (Table 4). Thus, individuals with higher initial values per individual brain outcome displayed less change compared to those with lower initial values per brain measure. Across brain regions, we observed the largest negative start-change correlations in gray matter directional intracellular diffusion and the smallest start-change correlations in macrostructure development (e.g., changes in GM volume, cortical area). Correlations between initial value and APΔ for measures of functional neurodevelopment and other aspects of brain microstructural development fell between these two relative extrema, with larger negative correlations in transverse diffusivity and functional connectivity, and smaller negative correlations, in gray-to-white matter contrast and BOLD variance.

**Table 4:** Associations between values at ages 9-10 years and annualized percent change across brain regions, per measure

| Measure | Partial correlation* |  |  |  |
| --- | --- | --- | --- | --- |
| | $r_{base,AP\Delta}$ | [Q<sub>1</sub>, Q<sub>3</sub>] | $r_{age,AP\Delta}$ | [Q<sub>1</sub>, Q<sub>3</sub>] |
| Macrostructure |  |  |  |  |
| GM volume | -0.16 | (-0.2, -0.1) | -0.05 | (-0.08, -0.02) |
| Cortical thickness | -0.33 | (-0.36, -0.26) | -0.04 | (-0.05, -0.02) |
| Cortical area | -0.16 | (-0.20, -0.11) | -0.05 | (-0.08, -0.03) |
| WM volume | -0.21 | (-0.25, -0.18) | 0.01 | (0.00, 0.03) |
| Microstructure |  |  |  |  |
| GM/WM contrast | -0.30 | (-0.35, -0.27) | -0.05 | (-0.06, -0.04) |
| Isotropic intracellular diffusion (GM) | -0.42 | (-0.48, -0.36) | 0.02 | (0.00, 0.05) |
| Directional intracellular diffusion (GM) | -0.64 | (-0.74, -0.58) | 0.00 | (-0.02, 0.03) |
| Fractional anisotropy (WM) | -0.40 | (-0.42, -0.36) | 0.03 | (0.02, 0.04) |
| Mean diffusivity (WM) | -0.48 | (-0.51, -0.43) | -0.02 | (-0.04, -0.01) |
| Transverse diffusivity (WM) | -0.29 | (-0.33, -0.24) | -0.02 | (-0.04, 0.01) |
| Longitudinal diffusivity (WM) | -0.52 | (-0.55, -0.49) | 0.01 | (0.00, 0.02) |
| Isotropic intracellular diffusion (WM) | -0.34 | (-0.40, -0.30) | 0.00 | (-0.02, 0.02) |
| Directional intracellular diffusion (WM) | -0.36 | (-0.38, -0.34) | 0.03 | (0.03, 0.04) |
| Function |  |  |  |  |
| BOLD variance | -0.33 | (-0.41, -0.33) | 0.00 | (-0.02, 0.02) |
| Within FC | -0.52 | (-0.53, -0.47) | 0.00 | (-0.01, 0.02) |
| Between FC | -0.52 | (-0.47, -0.44) | 0.01 | (0.00, 0.01) |
| Subcortical FC | -0.56 | (-0.58, -0.53) | 0.00 | (-0.02, 0.01) |

*Note.* \*Partial correlation between starting point and change (i.e., $r_{base,AP\Delta}$ ) removes the effect of age, while partial correlation between age and change (i.e., $r_{age,AP\Delta}$ ) removes the effect of starting point (i.e., values at ages 9-10 years). All partial correlations between starting point and change, for all regions, were significant at $\alpha_{corr} < 0.05$ . Abbreviations: GM = gray matter; WM = white matter; FC = functional connectivity; BOLD = blood-oxygen-level-dependent.

On the other hand, partial correlations between intra-individual change and age at study enrollment (i.e., 9-10 years) included both positive and negative associations, depending on the measure. Thus, older individuals demonstrated smaller brain changes than their younger counterparts on some measures, but larger brain changes on others. For example, cortical volume and thickness changes were negatively correlated with age at study enrollment, such that older participants demonstrated larger decreases in cortical volume and thickness over the following two years, while younger participants demonstrated smaller decreases. On the other hand, white matter volume changes were positively correlated with age at study enrollment, such that older participants demonstrated larger increases in white matter volume over the following two years, while younger participants demonstrated smaller increases. Similarly, older participants demonstrated larger increases in fractional anisotropy than did younger participants, but demonstrated larger decreases in mean diffusivity than did younger participants. White matter fractional anisotropy, longitudinal diffusivity, and gray matter isotropic intracellular diffusion all demonstrated positive age-change associations, while mean diffusivity, gray-to-white matter contrast, and transverse diffusivity all demonstrated negative age-change associations. Functional measures’ mean age-change associations were near or equal to zero, with interquartile ranges spanning zero to indicate that individual connections and regions included positive and negative age-change associations.

### 3.2. Inter-individual changes in brain development

Inter-individual differences of APΔ showed that brain changes were more similar between individuals in macro- and microstructural measures (Figure 3A), compared to functional measures, which were more variable between individuals. (Figure 3B).

**Figure 3.**
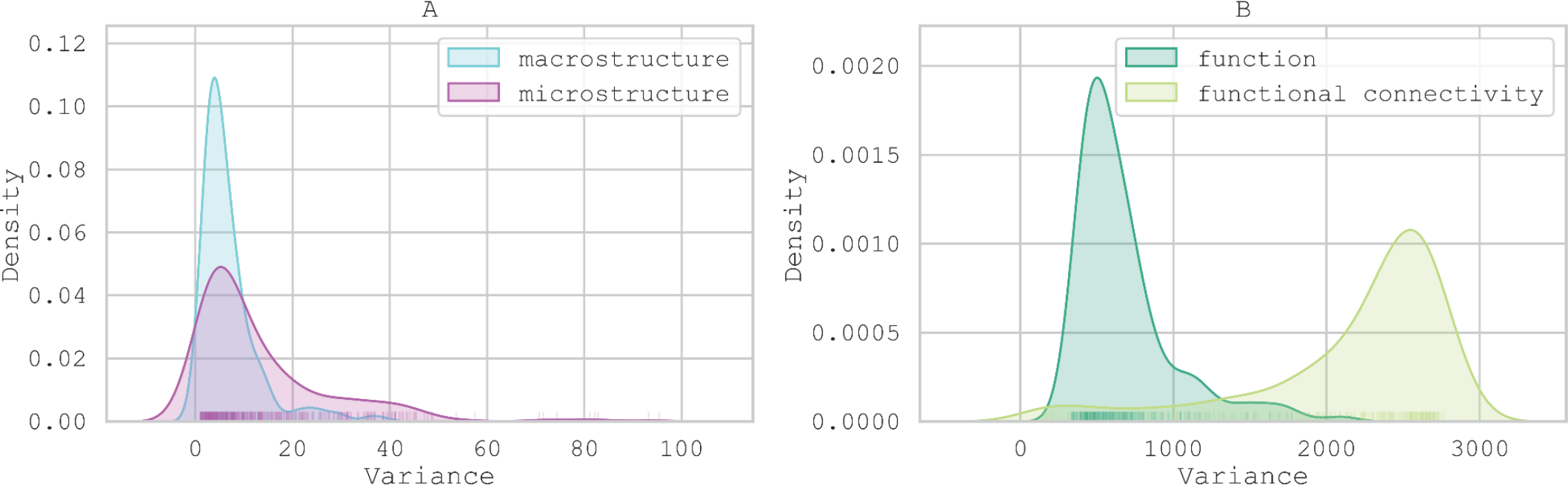
Distributions of inter-individual variance of annualized percent change in brain measures across all participants, separated by modality. For descriptions of the individual brain measures contributing to each broad category, see Table 2.

Comparisons of variance in APΔ across age bins, sex at birth, and pubertal status revealed significant heteroscedasticity across neuroimaging modalities, with inter-individual variability in the rate of brain macrostructure, microstructure, and functional development (Table 5). Figure 4 displays Fligner-Killeen (F-K) statistics,representing the amount of inhomogeneity between levels of each variable (i.e., age, sex, puberty) for each brain outcome examined per imaging measure. showing how unevenly individual differences in developmental change are distributed. Higher F-K statistics indicate greater differences in variance, or in other words greater inhomogeneity, in developmental change between age groups, sexes, and pubertal stages, respectively. For example, microstructure measures demonstrate less heterogeneity in inter-individual variance, or more equal distributions of individual differences, across participants of different ages at study enrollment, compared to macrostructure and functional measures. In contrast to the correlations between age and APΔ described above, heteroscedasticity tells us that there is greater variability between individuals of a certain age group at study enrollment as compared to others who enrolled in the study at an earlier or later age. Additional plots of variance seen for each neuroimaging measure by age, sex, and puberty categories are included in Supplemental Figures 2-4.

**Table 5:**
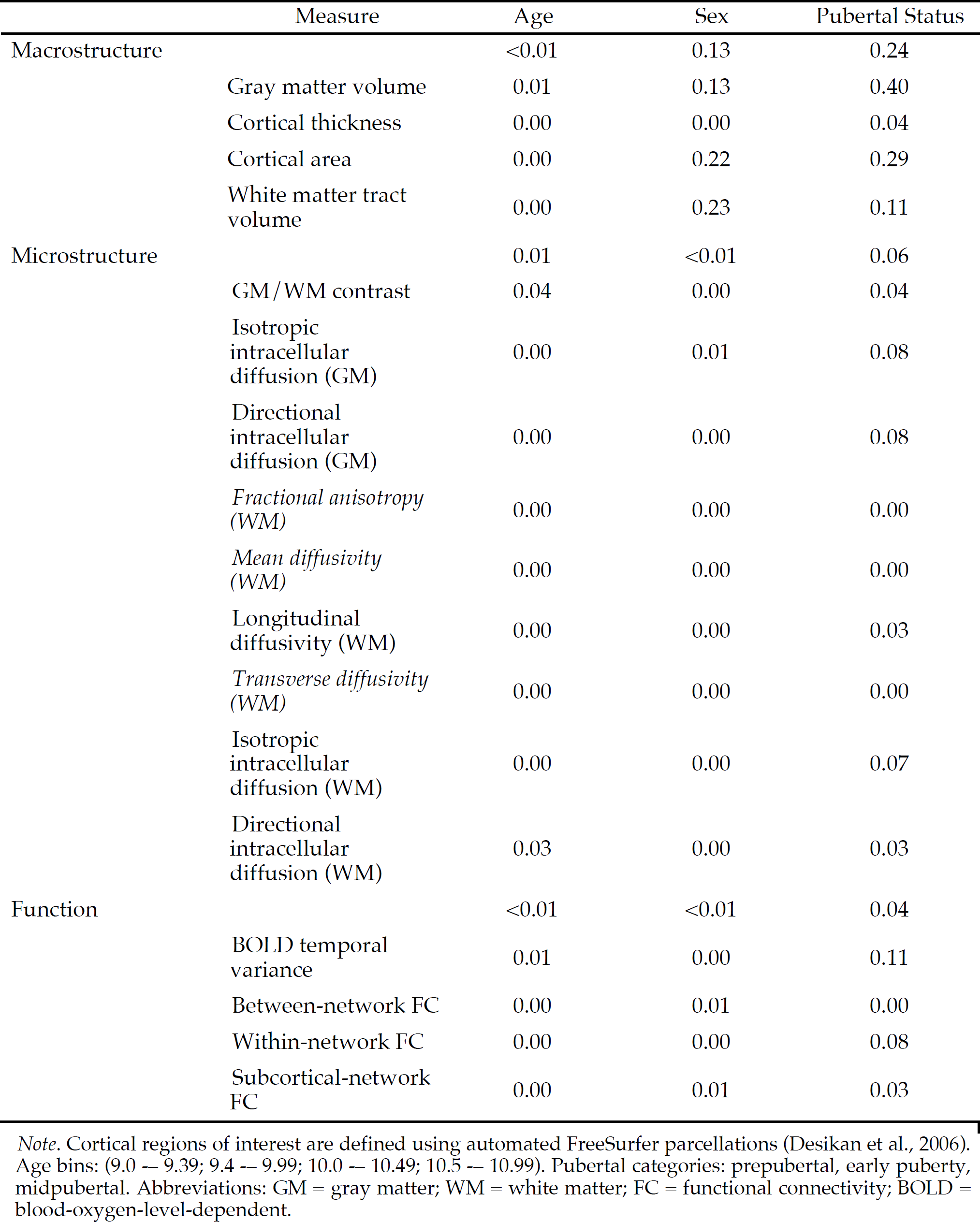
Proportion of brain regions that are significantly heteroscedastic for APΔ with respect to age, sex, and pubertal status at study enrollment

|  | Measure | Age | Sex | Pubertal Status |
| --- | --- | --- | --- | --- |
| Macrostructure |  | <0.01 | 0.13 | 0.24 |
|  | Gray matter volume | 0.01 | 0.13 | 0.40 |
|  | Cortical thickness | 0.00 | 0.00 | 0.04 |
|  | Cortical area | 0.00 | 0.22 | 0.29 |
|  | White matter tract volume | 0.00 | 0.23 | 0.11 |
| Microstructure |  | 0.01 | <0.01 | 0.06 |
|  | GM/WM contrast | 0.04 | 0.00 | 0.04 |
|  | Isotropic intracellular diffusion (GM) | 0.00 | 0.01 | 0.08 |
|  | Directional intracellular diffusion (GM) | 0.00 | 0.00 | 0.08 |
|  | Fractional anisotropy (WM) | 0.00 | 0.00 | 0.00 |
|  | Mean diffusivity (WM) | 0.00 | 0.00 | 0.00 |
|  | Longitudinal diffusivity (WM) | 0.00 | 0.00 | 0.03 |
|  | Transverse diffusivity (WM) | 0.00 | 0.00 | 0.00 |
|  | Isotropic intracellular diffusion (WM) | 0.00 | 0.00 | 0.07 |
|  | Directional intracellular diffusion (WM) | 0.03 | 0.00 | 0.03 |
| Function |  | <0.01 | <0.01 | 0.04 |
|  | BOLD temporal variance | 0.01 | 0.00 | 0.11 |
|  | Between-network FC | 0.00 | 0.01 | 0.00 |
|  | Within-network FC | 0.00 | 0.00 | 0.08 |
|  | Subcortical-network FC | 0.00 | 0.01 | 0.03 |

*Note.* Cortical regions of interest are defined using automated FreeSurfer parcellations (Desikan et al., 2006). Age bins: (9.0 — 9.39; 9.4 — 9.99; 10.0 — 10.49; 10.5 — 10.99). Pubertal categories: prepubertal, early puberty, midpubertal. Abbreviations: GM = gray matter; WM = white matter; FC = functional connectivity; BOLD = blood-oxygen-level-dependent.

**Figure 4.**
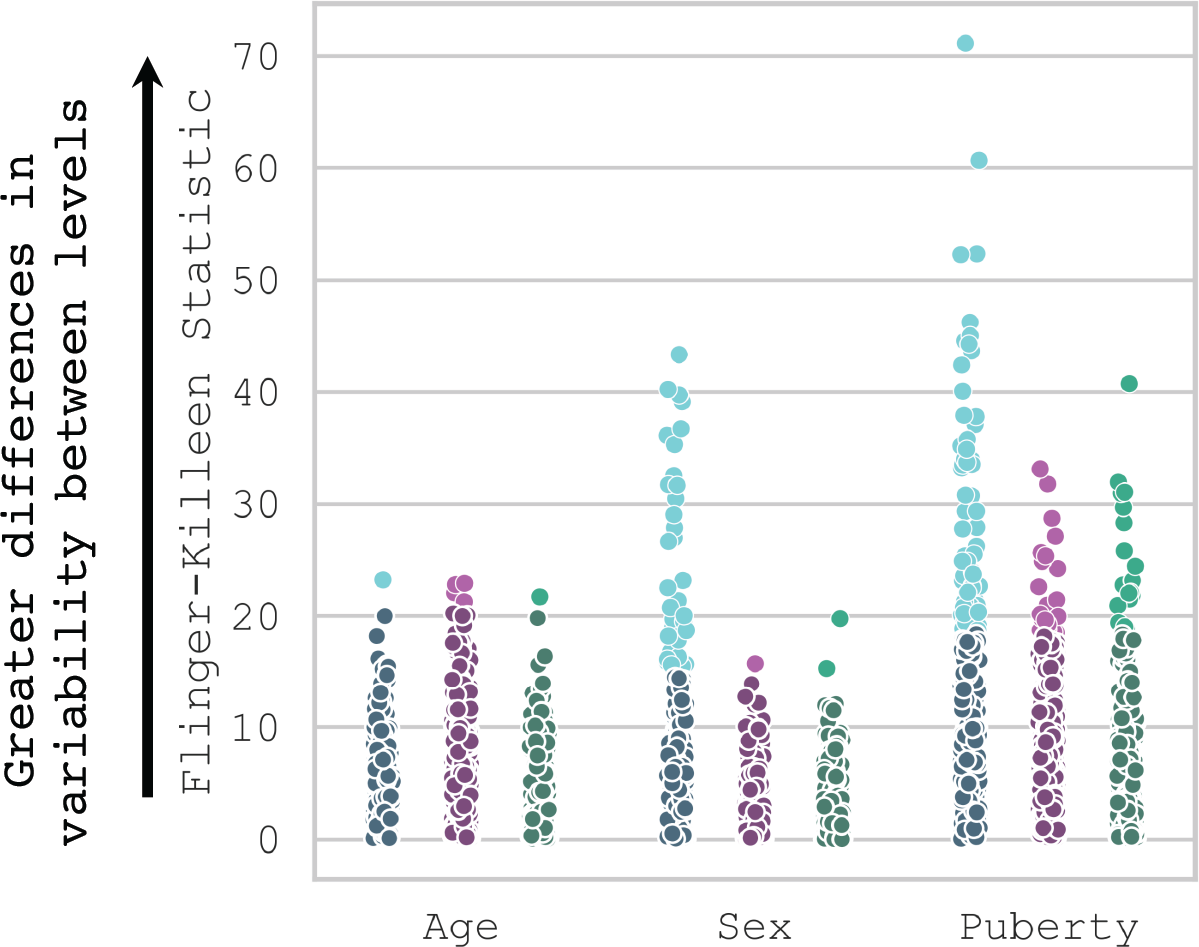
Magnitude and significance of heteroscedasticity in annualized percent change for each brain region by neuroimaging modality (per Fligner-Killeen, F-K, statistic) as a function of age (9.0 -– 9.39; 9.4 -– 9.99; 10.0 -– 10.49; 10.5 -– 10.99 years), sex (male, female), and pubertal status (pre, early, mid) at study enrollment. Significant heteroscedasticity of each brain region is indicated by a lighter marker; non-significant, by a darker marker. Color indicates neuroimaging modality (i.e., macrostructure, microstructure, function), each point represents heteroscedasticity of a single measure (e.g., brain region, global estimate). F-K statistics follow a *X*^2^ distribution, such that higher values indicate unequal variance between age, sex, and pubertal status categories.

Overall, most brain measures displayed similar variance in APΔ, or evenly distributed individual differences, based on initial age at study enrollment (binned in equal-sized increments from 9.0-10.99 years) as shown by low F-K scores. Less than or equal to 1% of regions or tracts demonstrated significant differences in variance across children based on their age at study enrollment across macrostructural (Figure 5A), microstructural (Figure 6A), and functional (Figure 7A) variance. With respect to sex, macrostructure changes demonstrated considerably more heteroscedasticity (∼13% of regions; Figure 5B), as compared to microstructure(<1% of regions or tracts; Figure 6B), or functional changes (<1% of regions or networks; Figure 7B). Unequal distribution of variance in APΔ was more prominent between individuals classified as being in prepubertal, early puberty, and midpubertal stages at study enrollment. With respect to puberty, 24% of regions or tracts across macrostructural measures (Figure 5C), 6% of GM and WM microstructural measures (Figure 6C), and 4% of functional measures (Figure 7C) demonstrated significant heteroscedasticity. While the majority of measures that were heteroscedastic across pubertal stages demonstrated the greatest variance in mid-puberty as compared to early and pre-pubertal development, white matter volume and longitudinal diffusivity demonstrated the greatest variance in pre-pubertal individuals as compared to early and mid-pubertal participants (Supplemental Figure 4).

**Figure 5.**
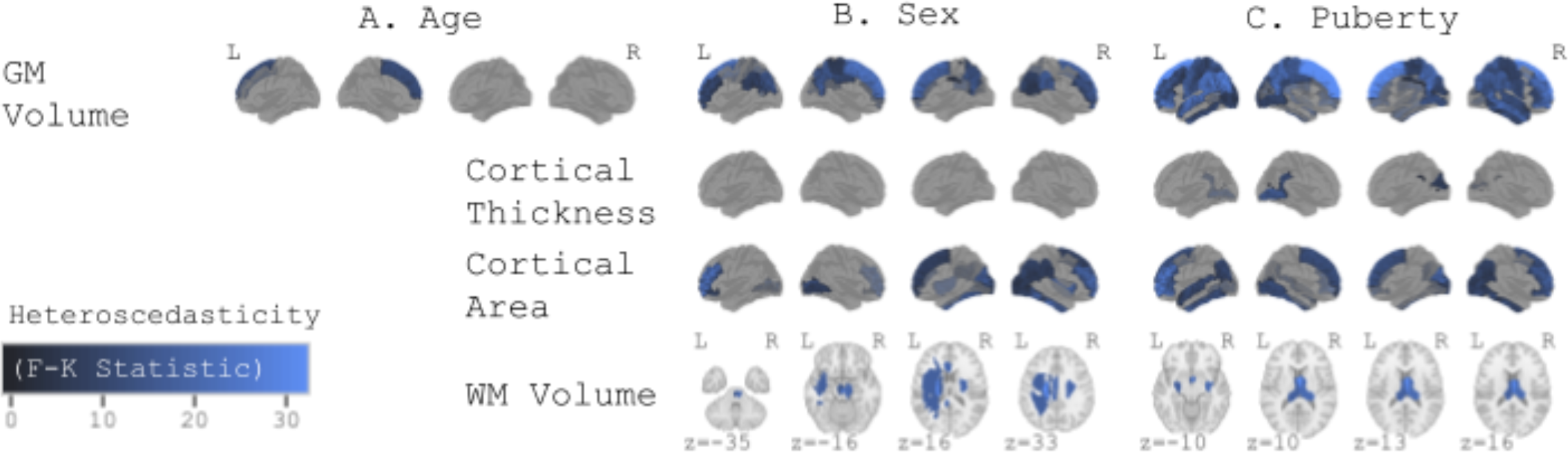
Heteroscedasticity of annualized percent changes in macrostructural brain development. Macrostructural measures of gray and white matter brain changes are represented in rows, while developmental variables (i.e., age, sex, puberty) are represented in columns. Brighter hues indicate greater heteroscedasticity of macrostructural brain development with respect to A) age (9.0 -– 9.39; 9.4 -– 9.99; 10.0 -– 10.49; 10.5 -– 10.99 years), B) sex (male, female), and C) pubertal stage (prepubertal, early puberty, mid-pubertal) at the time of study enrollment (i.e., larger differences in variance between levels). Abbreviations: GM = gray matter; WM = white matter; L = left; R = right; F-K = Fligner-Killeen.

**Figure 6.**
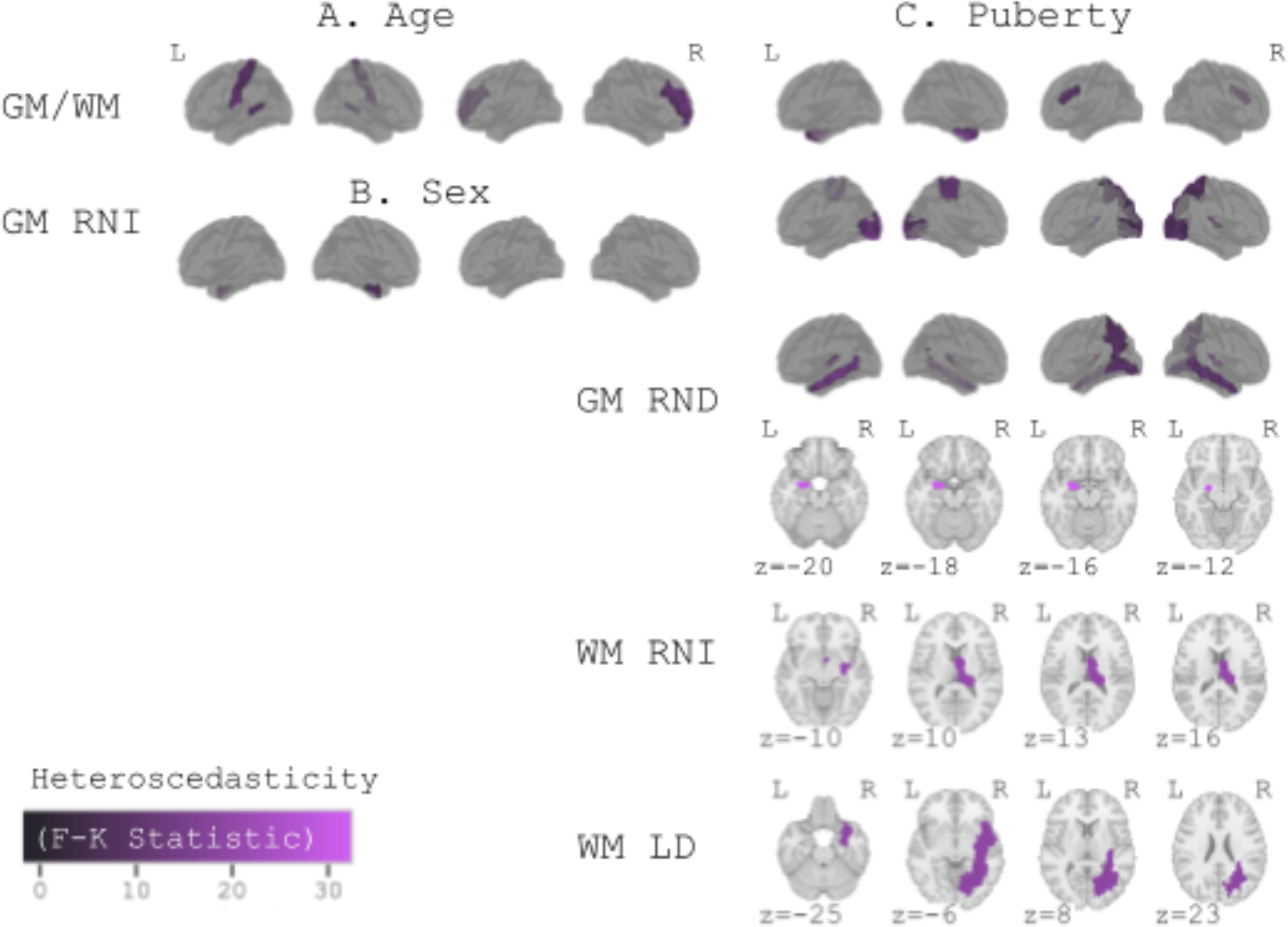
Heteroscedasticity of annualized percent changes in microstructural brain development. Microstructural measures of gray and white matter brain changes are represented in rows, while developmental variables (i.e., age, sex, puberty) are represented in columns. Brighter hues indicate greater heteroscedasticity of microstructural brain change with respect to age (9.0 -– 9.39; 9.4 -– 9.99; 10.0 -– 10.49; 10.5 -– 10.99 years), sex (male, female), and pubertal stage (prepubertal, early puberty, mid-pubertal) at the time of study enrollment (i.e., larger differences in variance between levels). Abbreviations: GM/WM = gray-to-white matter contrast; GM RNI = gray matter isotropic intracellular diffusion; GM RND = gray matter directional intracellular diffusion; WM RNI = white matter isotropic intracellular diffusion; WM LD = white matter longitudinal diffusivity; L = left; R = right; F-K = Fligner-Killeen.

**Figure 7.**
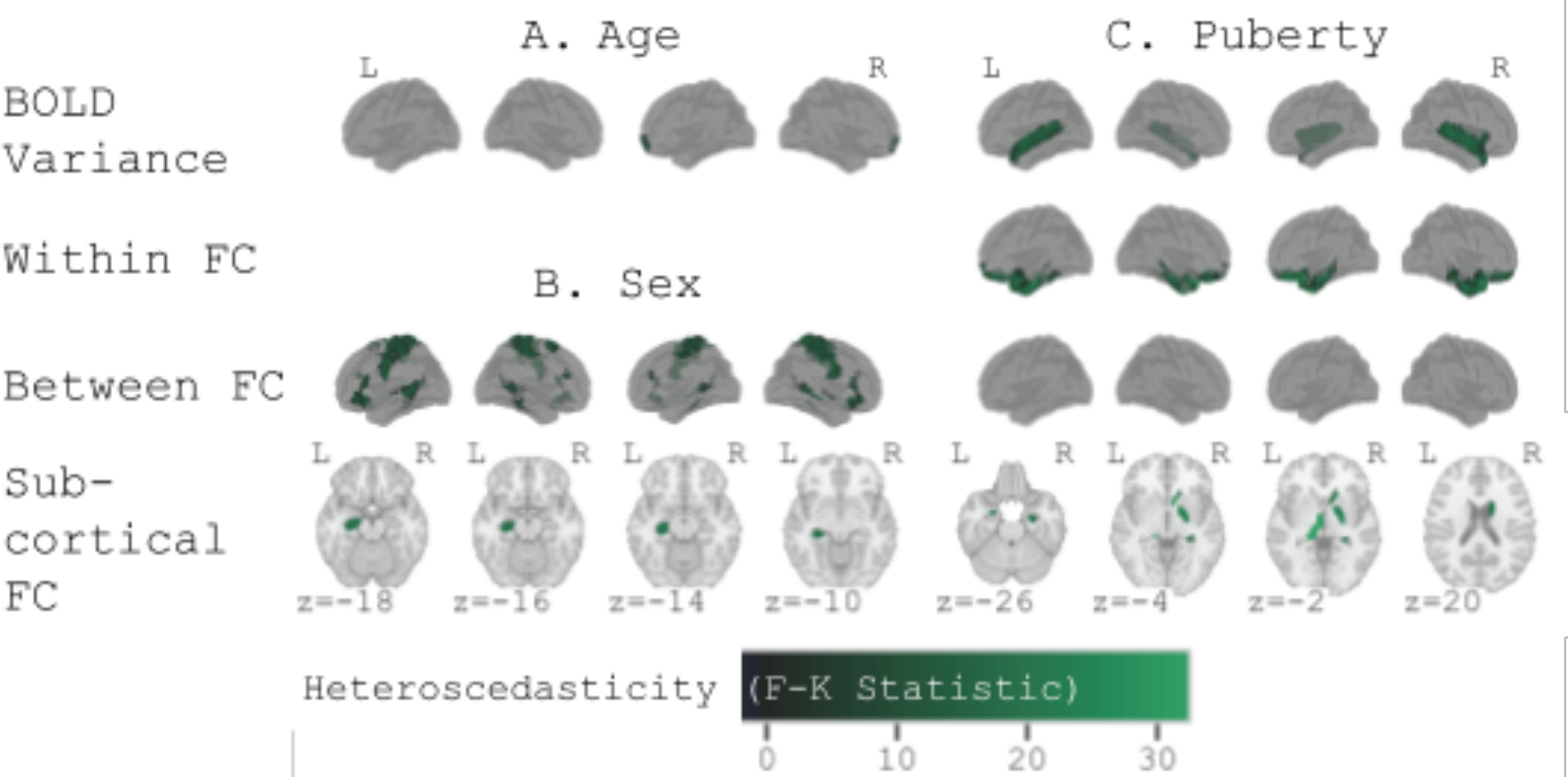
Heteroscedasticity of annualized percent changes in functional brain development. Functional measures of brain changes are represented in rows, while developmental variables (i.e., age, sex, puberty) are represented in columns. Brighter hues indicate greater heteroscedasticity of functional brain change with respect to A) age (9.0 -– 9.39; 9.4 -– 9.99; 10.0 - – 10.49; 10.5 -– 10.99 years), B) sex (male, female), and C) pubertal stage (prepubertal, early puberty, mid-pubertal) at the time of study enrollment (i.e., larger differences in variance between levels). Abbreviations: FC = functional connectivity; L = left; R = right; BOLD = blood-oxygen-level-dependent; F-K = Fligner-Killeen.

#### 3.2.1. Neuroanatomy of inter-individual changes in APΔ by age, sex, and pubertal status

Neuroanatomical differences in significantly heteroscedastic regions with respect to age, sex, and pubertal status at enrollment indicate that unique individual differences exist in respect to each. Below, we briefly describe these differences in neurodevelopment across neuroimaging measures of macrostructure, microstructure, and brain function.

##### 3.2.1.1. Macrostructure changes in neurodevelopment

Macrostructure APΔs exhibited differential variability with respect to age at study enrollment only in the medial superior frontal gyrus (Figure 5A). Sex- and puberty-related differences in variability of macrostructure changes were more widespread than those of age (Figure 5B, C). For example, greater APΔ variance was observed in the volumes of dorsal and midline cortical regions in female versus male children, while white matter volume in the corticospinal tract, cingulum, and superior longitudinal fasciculus demonstrated greater APΔ variance in male children than in female (Figure 5B, Supplemental Figure 3). Additionally, variance in the APΔ of lateral, posterior, and temporal GM volumes was significantly different across pubertal stages. Differences in APΔ white matter volume variability between pubertal stages was restricted to the fornix and corpus callosum (Figure 5C). Within regions of heteroscedastic cortical area and volume changes, pre-pubertal individuals demonstrated the greatest variability; mid-pubertal, the least; while the opposite was true for white matter volume changes (Supplemental Figure 4).

##### 3.2.1.2. Microstructure changes in neurodevelopment

Differences in variability of APΔ in microstructure across age and pubertal stage were far more limited compared to macrostructure and brain function.). Microstructural changes demonstrated differing variability in APΔ with respect to age only in gray-to-white matter contrast, which was primarily seen in postcentral, middle temporal, and middle frontal gyri (Figure 6A). Moreover, variability in APΔ of gray and white matter microstructure was only significantly different between males and females for gray matter isotropic intracellular diffusion of the entorhinal cortex (Figure 6B). Differences in APΔ variability between pubertal stages were most prominent for longitudinal diffusivity in the right corticospinal tract and inferior longitudinal fasciculus, as well as transverse diffusivity of the left uncinate fasciculus (Figure 6B). Similarly, significant heteroscedasticity in APΔ was seen in intracellular diffusion across pubertal stages, including both isotropic and directional intracellular diffusion, in posterior, temporal, and corticostriate gray matter regions (Figure 6C). Across microstructure changes that were heteroscedastic with respect to puberty, midpubertal children were the most variable for all measures except longitudinal diffusivity, in which prepubertal children were the most variable (Supplemental Figure 4).

##### 3.2.1.3. Functional changes in neurodevelopment

Individual differences in APΔ were more heterogeneously distributed across pubertal stages than between age bins or sexes (Figure 7A-C). Age and sex related differences in APΔ variability were most prominent for between-network functional connectivity, with much less heteroscedasticity seen across brain regions of interest for BOLD variance or within-network connectivity (Figure 7A,B). Changes in cingulo-opercular network functional connectivity to both default mode and dorsal attention networks exhibited differential variability with respect to age, as did connectivity of the putamen to extra-network regions (Figure 7A). Of functional changes, only connectivity between Retrosplenial Temporal and Ventral Attention Networks, and connectivity between Dorsal Attention Network and hippocampus demonstrated sex-related heteroscedasticity (Figure 7B). Changes in BOLD variance of bilateral superior temporal and right precentral gyri, in addition to functional connectivity of non-network cortical regions (Figure 7C). Within regions of heteroscedastic BOLD variance, pre-pubertal individuals demonstrated the least variability (Supplemental Figure 4).

#### 3.2.2. Inter-individual changes in APΔ as a function of pubertal status by sex and over time

We further assessed heteroscedasticity across pubertal stages at each wave of data collection (Supplemental Figure 5), and heteroscedasticity across changes in pubertal stage (Supplemental Figure 6 & 7); for (a) all participants, (b) female participants, and (c) male participants. Assessing within-sex heteroscedasticity with respect to puberty uncovered differing APΔ variability across pubertal stages in female and male participants. In female participants, macrostructural, microstructural, and functional measures demonstrated limited heteroscedasticity at ages 9-10 years, while at ages 10-11 years, only functional measures were heteroscedastic across pubertal stages. In contrast, no heterogeneity in APΔ variance was observed in any outcome measure across pubertal stages in female participants at ages 11-12 years. At ages 9-10 years in male participants, only macrostructural and functional measures were heteroscedastic across pubertal stages. In males at ages 10-11 and 11-12 years, only functional measures remained heteroscedastic across pubertal stages. Furthermore, no measures demonstrated heterogeneity in APΔ variance related to *changes* in pubertal stage across waves of data collection. Thus, microstructural heteroscedasticity across pubertal stages at ages 9-10 is driven by female participants, and differences in how individuals change as a function of puberty is largely based on the “starting point” (i.e., ages 9-10 years) in both male and female participants. The lack of heteroscedasticity with respect to changes in participants’ pubertal stages between ages 9-10 and 11-12 years further highlights that an individual’s “starting point” in puberty, in addition to brain measures (see section 3.1.2.), is important to individual differences in neurodevelopment across this age range.

## 4. Discussion

Here, we assessed within- and between-individual variability in brain development between ages 9-13 years. We characterized annualized percent change in macrostructural, microstructural, and functional measures in a large sample from the longitudinal ABCD Study, and then compared these rates of change across age, sex, and pubertal stage at study enrollment. These analyses revealed varying magnitudes of differences in individual variability across brain regions, tissue types, and facets of development. There are instances in which intra-individual variance in developmental changes are not equally distributed across sexes, ages, or pubertal stages; albeit this heterogeneity in inter-individual differences, varies across brain regions and measures of brain macrostructural, microstructural, and functional change. This work represents an important contribution to the study of individual differences in child brain development, which has long focused on mean trajectories and mean differences, by providing much-needed characterizations of within- and between-individual variance.

### 4.1. Moving away from the mean: Widespread individual differences in brain changes from 9-13 years of age

The existing literature on human brain development relies largely on cross-sectional or sequential cohort designs, characterizes development across a much broader age range, and/or focuses on normative or group-level changes over time. Comparatively little focus has been on *variability* in change. Our findings highlight that even within the narrow developmental window of early adolescence, intra-individual change is vastly noticeable and varies considerably from person to person. On a whole-brain scale, we identified average annualized percent changes (APΔ) consistent with literature on normative brain development across this age range. These include decreases, on average, in gray matter volume (Bethlehem et al., 2022; Lenroot & Giedd, 2006), cortical thickness (Tamnes et al., 2017; Wierenga et al., 2014), gray-to-white matter contrast (Norbom et al., 2019; Paus et al., 2001), mean diffusivity (Schmithorst & Yuan, 2010), and transverse diffusivity (Asato et al., 2010). We additionally replicated increases in white matter volume, fractional anisotropy (Lebel & Deoni, 2018; Schmithorst & Yuan, 2010; Tamnes et al., 2018), white and gray matter isotropic intracellular diffusion, white matter directional intracellular diffusion (Palmer et al., 2022), and within-network functional connectivity (Fair et al., 2007; Grayson & Fair, 2017; Satterthwaite et al., 2012). Also consistent with prior findings, we identified both increases and decreases in gray matter directional intracellular diffusion (Palmer et al., 2022), subcortical-network functional connectivity (Ji et al., 2019; Langen et al., 2018; van Duijvenvoorde et al., 2019), and BOLD variance (Nomi et al., 2017; Wang et al., 2021), depending on the brain region or network. Larger decreases in gray matter volume were identified in parietal regions than in frontal, temporal, or occipital regions, consistent with prior findings (Lenroot et al., 2007). Fine-grained comparisons with prior literature across microstructural measures are more difficult, unfortunately, as many studies of white matter microstructural development capture a much broader age range (e.g., 5 to 30 years), across which trajectories follow curvilinear trajectories, and studies of gray matter microstructural development are uncommon. However, there are a few notable similarities, including greater annualized percent change in fractional anisotropy of the cingulum than other tracts, along with virtually no change in the fornix (Lebel et al., 2008). Across functional measures, regional and network-wise comparisons with prior literature are difficult, too, given the narrow age range and overall lack of consensus across functional imaging studies of development (Oldham & Fornito, 2019), and differences in large-scale network definitions across the literature (Uddin et al., 2019).

On the other hand, the current estimates of developmental change conflicted with some prior findings. Compared with Sowell et al. (Sowell et al., 2004), this work only found increased cortical thickness in a much more restricted area, the bilateral entorhinal cortex, instead of extended temporal pole and orbitofrontal regions. However, our age range was a bit older than that of the sample in that study. In contrast to integration within networks, findings regarding segregation between large-scale functional brain networks were contrary to prior literature. We found that each network’s changes in connectivity to other networks were zero on average, indicating that changes in functional connectivity strength occur on a connection level, including weakened connectivity with some networks and strengthened connectivity with others, rather than as broad network-level segregation. Finally, there is not much literature on the development of intracellular diffusion across this age range. Here, gray matter directional intracellular diffusion showed opposite directions of change between cortical and some subcortical regions, potentially reflecting differential cytoarchitectonic processes underlying the development of cortical and subcortical regions. While this work moves toward a more comprehensive understanding of developmental change in brain structure and function in late childhood and early adolescence, further time points will be crucial to clarifying these different trajectories.

This work also extends recent findings demonstrating that individual rates of change (i.e., APΔ) are significantly associated with their starting point (here, values at ages 9-10 years) (Mills et al., 2021). Our current study not only extends prior findings into a larger and more generalizable sample of children, it provides evidence that this association generalizes to all brain measures included here; although to varying degrees across brain regions. That is, after controlling for age effects, changes in all brain regions were negatively associated with their initial values at ages 9-10 years, across all measures of brain macrostructure, microstructure, and function. It is worth noting that the “starting points” in this research refer only to the start of *measurement* in a particular study. A wide range of influences contribute to individual differences in brain development, including genetics and gene-by-environment interactions, as well as socioeconomic status and maternal experiences during pregnancy (Blokland et al., 2012; Brito & Noble, 2014; Farah, 2017; Gao et al., 2015; Gilmore et al., 2018; Harden et al., 2007; Turkheimer et al., 2003; Ursache & Noble, 2016). Such factors can also influence the pace of development (Tooley et al., 2021) and are, thus, likely reflected in the overall changes in brain development reported here. Brain development is further complicated by changes in the relative influence of genes versus the environment throughout childhood and adolescence (Lenroot & Giedd, 2006).

Our findings may reflect curvilinear intra-individual developmental trajectories from 9-12 years, such that higher values at ages 9-10 years are associated with less positive change in the next two years. This is consistent with much of the group-level effects reported in the literature on normative trajectories in structural brain development (Bethlehem et al., 2022; Mills et al., 2016; Wierenga et al., 2014), although there is some evidence of linear developmental trajectories, for example in gray-to-white matter contrast and in white matter area (Norbom et al., 2019; Paus et al., 2001; Wierenga et al., 2014). Finally, these associations may reflect *equipotentiality* as presented in developmental psychobiology, which implies that many paths in development can lead to the same outcomes or consequences (Bornstein, 2018). For example, while the rate may differ between individuals, the sequence of development is similar, so that individuals with lower brain metric values at ages 9-10 years had either more increasing or less decreasing to do towards neurodevelopmental milestones, depending on the neuroimaging measure and brain region.

### 4.2. Inter-individual variability in brain maturation is not equally distributed across imaging measures

Recently, more research has focused on leveraging large datasets to study individual differences in development, but these studies still focus largely on differences in mean trajectories. Contextualizing mean trajectories by assessing developmental variance is necessary to comprehensively understand human brain development. Characterizing developmental variance illuminates the ability of normative trajectories to describe individual-level phenomena. Across this sample, functional changes were much more variable than were changes in macrostructure and microstructure. Within measures of macrostructural change, white and gray matter volume showed the greatest inter-individual variability, whereas heterogeneity in annualized change in cortical thickness was minimal. In general, measures of microstructural change showed more inter-individual variability than did macrostructural measures, with the greatest variability seen in directional intracellular diffusion in gray matter. Inter-individual variability in functional measures was an order of magnitude greater than in structural measures, with subcortical-cortical network connectivity showing the greatest variability and within-network connectivity, the least. With the current MRI data it is difficult to disentangle variability inherent to the MRI scanner and scanning sequence from variability due to underlying neurobiological differences between individuals, though this certainly warrants further study. Differences in variability between macrostructural, microstructural, and functional measures may be due to differences in neurobiological variability or they may be due to differences in image acquisition, processing, and inherent measurement error to a given modality (Birn et al., 2013; Botvinik-Nezer et al., 2020; Kirilina et al., 2016).

### 4.3. Inter-individual variability: Brain changes vary between individuals in early adolescents

This work represents a novel application of a classic statistical concept, using heteroscedasticity to assess distributions of neurodevelopmental variability across age, sex, and puberty in children ages 9-13 years. Thus, we describe not only the variability in neurodevelopmental change, but its distribution across developmental factors. We identified few age-related differences in inter-individual variability (i.e., heteroscedasticity) across measures, including changes in gray matter volume, white matter isotropic intracellular diffusion, and BOLD variance. Interestingly, variability in gray matter volume increased with increasing age, while variability in the other measures demonstrating significant heteroscedasticity across age showed the greatest variability in individuals ages 9.5-10 years at the beginning of data collection. We expect these nuanced differences in distributions of inter-individual variability, within both measures of brain change *and* specific age ranges, likely reflecting the differential timing and neuroanatomy of aspects of brain development in this age range.

Sex differences in variability of brain macrostructure have been previously noted across the lifespan, dominated by greater variability in males compared to females (Wierenga et al., 2022), often referred to as the greater male variability hypothesis. However, this work has been largely cross-sectional, with less information concerning variability in change. Our findings address that gap and show heteroscedasticity in cortical volume, area, and intracellular diffusion; white matter volume; and functional network connectivity exhibit sex differences in *variability* as it pertains to developmental change. However, these findings only support the greater male variability hypothesis for a few measures of brain change. Male children exhibited greater variability in tracts with significantly heteroscedastic white matter volume, and in heteroscedastic network connectivity (both cortical and subcortical). On the other hand, female children showed greater variability in regions of significantly heteroscedastic cortical area, volume, and intracellular diffusion. Changes in most brain regions across each of these measures displayed no sex differences in variability, nor did most brain measures overall. Together, this work extends Wierenga and colleagues’ findings to suggest greater male-than-female variability in *changes in* structural (i.e., white matter) and functional connections exist, in addition to established cross-sectional differences, along with greater female-than-male variability in cortical macro- and microstructure changes during early adolescence. Thus, we provide some neurobiological limitations to the larger variability hypothesis that posits greater male-than-female variability across a range of psychological and physical attributes (Hyde, 2014; Johnson et al., 2008; Lehre et al., 2009; Winsor, 1927).

Puberty plays a role in neurodevelopmental trajectories (Dai & Scherf, 2019; Goddings et al., 2014; Herting, Johnson, et al., 2018; Vijayakumar et al., 2018), such that individual differences in hormones (Herting et al., 2014; Vijayakumar, Youssef, Allen, Anderson, Efron, Mundy, et al., 2021), pubertal tempo (Vijayakumar et al., 2018; Vijayakumar, Youssef, Allen, Anderson, Efron, Hazell, et al., 2021; Vijayakumar, Youssef, Allen, Anderson, Efron, Mundy, et al., 2021), and pubertal staging are associated with brain development (Goddings et al., 2019; Herting & Sowell, 2017). Here, we have identified differences in the variability of annualized brain changes across Tanner stages of pubertal development during the narrow age-range of ages 9-10 years. Interestingly, these changes are more evenly distributed (i.e., less heteroscedastic) across pubertal stages at ages 10-11 years and even more so at ages 11-12 years. Moreover, we show the patterns of variability in neurodevelopment seen across Tanner stages at ages 9-10 years are distinct from and more widespread than those seen across ages and between sexes. Globally, prepubertal individuals exhibit greater variability in heteroscedastic white matter changes (i.e., tract volume and mean diffusivity), while mid-pubertal individuals exhibit greater variability across cortical and subcortical heteroscedastic brain changes (cortical thickness, area, volume, and intracellular diffusion; BOLD variance, and functional connectivity). Taken together, the diversity of inter-individual variability in brain development suggests only some aspects of neurodevelopment display globally monotonic increasing or decreasing of variability as a function of puberty at ages 9-10 years. Further research is needed to better understand this diversity in neurodevelopmental trajectories with respect to puberty and whether these patterns reflect differential susceptibility of neurobiological mechanisms between brain regions.

### 4.4. The importance of heteroscedasticity to developmental cognitive neuroscience

The current study provides novel insight into intra-individual and inter-individual differences in brain development, with notable heterogeneity in annualized patterns of change across the narrow age range of 9-13 years-old. Understanding variability in brain changes and factors contributing to this variability across common developmental categories is essential. Large-scale, univariate studies have highlighted the importance of “unmodeled noise”, or the often overlooked sources of intra- and inter-individual variability that can confuse or obscure brain-phenotype associations (Bandettini et al., 2022; Dubois & Adolphs, 2016). Thus, teasing apart this unmodeled noise is crucial for robust study of brain-phenotype associations in development. Furthermore, profiling variability and sources thereof (e.g., via heteroscedasticity) is crucial for studying normative development and deviations therefrom, which can help illuminate the etiology of neurodevelopmental disorders and the emergence of psychopathology during this crucial period. Well-characterized normative models require robust descriptions of both central tendency *and* spread of the data. While developmental cognitive neuroscience has made recent, substantial advancements in describing average trajectories, the variation in these trajectories as highlighted here has received comparably little attention. Studying how deviations from normative development underlie neurodevelopmental disorders, the emergence of psychopathology, and social determinants of health depends on, first, understanding the scope and magnitude of normative variations in development. Profiling variation associated with age, sex, and puberty both across various brain regions and brain measures can provide information about which brain regions, tracts, and networks may be more susceptible to risk factors during adolescent development, and the neurobiological processes underlying this susceptibility. Finally, identifying developmental periods and groups of individuals that demonstrate greater inter-individual differences provides clear targets for further study. Thus, the context provided by this work has sizable utility for studying individual differences in brain development and the broad survey of imaging measures maximizes this utility for researchers with a range of interests and methodologies.

### 4.5. Limitations & Future Directions

Given that developmental trajectories can be better characterized by three or more timepoints, this line of investigation, assessing variability in brain trajectories, will continue to be important to investigate as additional ABCD Study data is released in future years.

Noise introduced by participant head motion is a nontrivial confound in resting state fMRI analyses and there are significant age, clinical, cognitive, and sociodemographic differences between participants exhibiting greater and less head motion in the ABCD cohort (Cosgrove et al., 2022). These differences limit the generalizability of resting state fMRI, confound studies that use these data, and may also contribute to some of the heteroscedasticity in functional measures identified here. Further, another factor that may confound estimates of variability between MRI modalities (i.e. macrostructure volumes, white matter tracts, and functional networks) is the size of brain regions, tracts, and networks. That is, larger regions represent average values for a greater number of data points (voxels) and, thus, may be more robust to both small, meaningful differences and random noise, resulting in less variability. On the other hand, within-region, -tract, or -network heterogeneity may increase variability of any given measure. Relatedly, it is impossible to remove the impact of differences in test-retest reliability between MRI sequences used to estimate macrostructural (i.e., T1-weighted scans), microstructural (i.e., diffusion-weighted scans), and functional (i.e., BOLD fMRI scans) on between-measure differences in variability. For example: although not included here, the task-based fMRI data from the ABCD Study demonstrate notably low reliability even within a session (Kennedy et al., 2022). On the other hand, between-individual differences have been shown to swamp scanner-induced variability across measures of macrostructure, microstructure, and function in an adult sample (Hawco et al., 2018).

Moving to population-level study brings new challenges to neuroimaging research, which has historically depended on small, homogeneous samples, as opposed to the large, diverse samples required for generalizable research. In this transition, researchers should consider the role that various sources of bias play in their work, such as those outlined in bias assessment tools commonly used in environmental epidemiology (Eick et al., 2020), and assess how the structure of variance is associated with sample characteristics and data collection.

## 5. Conclusions

Here, we provide a much-needed assessment of variability in intra-individual change during late child and early adolescent development, along with novel insight into heterogeneity in this variability across ages, sexes, and pubertal stages. Annualized percent change estimates suggest both intra-individual and inter-individual differences in trajectories of macrostructural, microstructural, and functional brain development throughout the brain from 9-13 years of age. These findings include novel insight into the magnitude of annual changes in both gray and white matter intracellular diffusion. Large-scale brain networks exhibited increased within-network connectivity overall, but, contrary to prevailing ideas about functional network segregation during development, connectivity between networks both increased and decreased, depending on the networks. Across most measures of brain macrostructure, microstructure, and function, individuals with smaller starting values displayed larger changes in brain development over the 2-year follow-up period. Functional measures exhibited much greater inter-individual variability than did structural measures, with changes in functional connectivity exhibiting the greatest variability. Individual differences in change were not equally distributed across pubertal stages and, to a lesser extent, ages and sexes, though these patterns differed with respect to imaging measure and brain regions. Assessments of homogeneity in variance across age, sex, and puberty revealed only limited support for the greater male variability hypothesis and for the hypothesis of greater variability between individuals in later stages of puberty. The current study represents important context and an insightful starting point for researchers who are interested in understanding individual differences in childhood and adolescent development.

## Acknowledgments

A special thank you to all of the children and families for their participation in their ABCD Study.

Research described in this article was supported by the National Institutes of Health [NIEHS R01ES032295] CCI would like to acknowledge scholars involved in NSP (R25 NS089462), BRAINS (R25 NS094094), and Diversifying CNS (R25 NS117356), as well as R25MH125545 and R25MH120869 for creating a supportive network of ABCD Study users.

Data used in the preparation of this article were obtained from the Adolescent Brain Cognitive DevelopmentSM (ABCD) Study (https://abcdstudy.org), held in the NIMH Data Archive (NDA). This is a multisite, longitudinal study designed to recruit more than 10,000 children age 9-10 and follow them over 10 years into early adulthood. The ABCD Study® is supported by the National Institutes of Health and additional federal partners under award numbers U01DA041048, U01DA050989, U01DA051016, U01DA041022, U01DA051018, U01DA051037, U01DA050987, U01DA041174, U01DA041106, U01DA041117, U01DA041028, U01DA041134, U01DA050988, U01DA051039, U01DA041156, U01DA041025, U01DA041120, U01DA051038, U01DA041148, U01DA041093, U01DA041089, U24DA041123, U24DA041147. A full list of supporters is available at https://abcdstudy.org/federal-partners.html. A listing of participating sites and a complete listing of the study investigators can be found at https://abcdstudy.org/consortium_members/. ABCD consortium investigators designed and implemented the study and/or provided data but did not necessarily participate in the analysis or writing of this report. This manuscript reflects the views of the authors and may not reflect the opinions or views of the NIH or ABCD consortium investigators. The ABCD data repository grows and changes over time. The ABCD data used in this report came from http://dx.doi.org/10.15154/1523041.

## Author Contributions

Conceptualization: KLB, MMH

Data curation: KLB

Formal Analysis: KLB

Funding acquisition: MMH

Methodology: KLB, MMH

Project administration: KLB, MMH

Resources: MMH

Software: KLB

Supervision: MMH

Visualization: KLB

Writing – original draft: KLB, MMH

Writing – review & editing: MMH, KLB, CCI, KLM, ARL

