## Supplemental Information for "Profiling intra- and inter-individual differences in brain development across early adolescence"

This document includes:

- Supplemental Methods
- Supplemental Results
- Supplemental Tables 0 to 4
- Supplemental Figures 1 to 7

#### Supplemental Methods

##### Summary of code used to perform these analyses and generate relevant figures, tables

All code used to perform the analyses reported here, and to generate related figures and tables, is available at github.com/62442katieb/deltaABCD_variability.

*Corresponding Author*: **Dr. Katherine L. Bottenhorn**, Postdoctoral scholar, Department of Population and Public Health Sciences, University of Southern California, Los Angeles, CA |

**Supplemental Table 0.** Code-to-analysis mapping

| Script | Purpose | Figures | Tables |
|---|---|---|---|
| `0.0datawrangling.py` | | | |
| | Extract relevant variables from .csv files downloaded from NDA and computed in-house (i.e., change scores); convert to wide format; assemble data dictionary | | |
| `0.1sample_demographics.py` | | | |
| | Calculate the number of individuals in each demographic group (both in whole ABCD dataset and in study-specific subset), re-binned for approximately equal group sizes to enable heteroscedasticity calculations | | Table 1, Supplemental Tables 1 & 2 |
| `0.2nifti_to_variable_mapping.py` | | | |
| | Create a dataframe that maps ABCD brain-related variable names to the associated nifti image files and values within those files, to enable plotting of AP$\Delta$ and heteroscedasticity values in brain space. | | |
| `1.0variance.py` | | | |
| | Calculate variance; rebin developmental and demographic variable (i.e., age, sex, pubertal status, scanner manufacturer, household income, race/ethnicity, caregiver education, & caregiver marital status); calculate and plot heteroscedasticity per brain measure, per variable. | Figure 3 | |
| `1.1parsing_heteroscedasticity.py` | | | |
| | Tabulate and plot heteroscedasticity of brain measures across developmental variables | Figures 4-7, Supplemental Figures 2-4 | Table 5 |
| `1.2change_score_descriptives.py` | | | |
| | Calculate and plot descriptive statistics for estimates of annualized percent change, per region/tract/network, per measure. | Figure 1 Figure 2 | Table 3 |
| `1.4start_change.py` | | | |
| | Compute Pearson's and partial correlations between baseline values, baseline age, and AP$\Delta$ across brain measures; plot results | | Table 4 |
| `1.5response_to_reviewers.py` | | | |
| | Compare pubertal stages between male and female participants, and across data collection time points (i.e., values from baseline at ages 9-10 years, one-year follow-up at ages 10-11 years, and two-year follow-up at ages 11-12 years) to disentangle roles of sex and puberty over time in inter-individual variability. | Supplemental Figures 4 - 7 | Supplemental Table 3 |

*Note*: These scripts include code to address heteroscedasticity of brain measures across demographic variables and scanner manufacturer that is not reported in this manuscript. They will be reported in a forthcoming manuscript.

**Supplemental Table 1.** Demographic Differences Across Scanners

| | Siemens | Philips | GE | $X^2$ | $p(X^2)$ |
|---|---|---|---|---|---|
| N (%) | 4539 (60%) | 905 (12%) | 2013 (28%) | | |
| ***Age (baseline)*** | 119.08 ± 7.38 | 118.89 ± 7.27 | 117.92 ± 7.59 | 4.45x10$^7$ | 0.04 |
| Sex Assigned at Birth | | | | 2.98 | 0.22 |
| Male | 2479 (55%) | 488 (54%) | 1053 (52%) | | |
| Female | 2060 (45%) | 417 (46%) | 960 (48%) | | |
| ***Race, Ethnicity*** | | | | 239.27 | 0.00 |
| Black | 694 (15%) | 109 (12%) | 178 (9%) | | |
| White | 2634 (58%) | 545 (60%) | 929 (46%) | | |
| Hispanic | 746 (16%) | 158 (17%) | 546 (27%) | | |
| Asian, other | 465 (10%) | 93 (10%) | 360 (18%) | | |
| ***Household Income*** | | | | 67.13 | 0.00 |
| >$100k | 1765 (39%) | 398 (44%) | 713 (35%) | | |
| $50 to $100k | 1340 (30%) | 243 (27%) | 499 (25% | | |
| <$50k | 1113 (25%) | 182 (20%) | 627 (31%) | | |
| **Caregiver Marital Status** | | | | 19.22 | 0.01 |
| Married | 3185 (70%) | 658 (73%) | 1349 (67%) | | |
| Widowed | 38 (<1%) | 6 (<1%) | 16 (1%) | | |
| Divorced | 381 (8%) | 76 (8%) | 202 (10%) | | |
| Separated | 145 (3%) | 28 (3%) | 89 (4%) | | |
| Never | 545 (12%) | 86 (10%) | 219 (11%) | | |
| Refused | 19 (<1%) | 6 (<1%) | 18 (1%) | | |
| ***Caregiver Education*** | | | | 72.36 | 0.00 |
| up to HS, GED | 650 (14%) | 136 (15%) | 361 (18%) | | |
| Some College or Associate's degree | 1422 (31%) | 200 (22%) | 611 (30%) | | |
| Bachelor's degree | 1400 (31%) | 265 (29%) | 539 (27%) | | |
| Graduate degree | 1060 (23%) | 303 (33%) | 500 (25%) | | |

*Note*. Omnibus tests for differences in distribution across each variable between scanner manufacturers were performed using Chi-square tests. After *N*, percentages reflect the proportion in each group within that scanner manufacturer. Bold categories indicate significant differences at $\alpha < 0.05$; bold and italicized categories, at $\alpha < 0.01$.

#### Imaging data acquisition

The details of ABCD Study neuroimaging data acquisition have been published elsewhere (Casey et al., 2018). Here, we only provide details relevant to this work.

Across 21 data acquisition sites, imaging data for the ABCD Study were collected on 3T MRI scanners from three manufacturers: Siemens (PRISMA models), General Electric (GE; 750 model), and Philips, all with multi-channel head coils compatible with multiband echo planar imaging (EPI) sequences. Scanning protocols were completed in either 1 or 2 sessions (same day) at visits every two years, from which we use baseline imaging data (collected in 2016 - 2018 while participants were 9-10 years old) and year-2 follow-up data (collected 2018 - 2020, while participants were 11-12 years old). Prior to scanning, participants completed motion compliance training in an MR-environment simulator.

Scanning sessions started with a 3D T1-weighted MPRAGE scan with prospective motion correction and a 256x256mm FOV; 208 axial slices; TR/TE = 2500/2.88ms (Siemens), 6.3/2.9ms (Philips), or 2500/2.0ms (GE); TI = 1060ms; an $8^o$ flip angle resulting in $1mm^3$ isotropic resolution. Then, participants completed two 5-minute resting-state functional scans (eyes open, with a fixation cross) using a blood-oxygen level-dependent (BOLD) EPI sequence with fast integrated distortion correction and a 216x216mm FOV, 60 interleaved axial slices, TR/TE = 800ms/30ms, a $52^o$ flip angle, mutliband factor = 6; in-plane GRAPPA acceleration resulting in $(2.4mm)^3$ isotropic resolution. Then, a T2-weighted structural scan was performed (not used here) followed by a high angular resolution diffusion imaging (HARDI) diffusion-weighted scan with a 240x240mm FOV; 81 axial slices; TR/TE = 4100/88ms (Siemens), 5300/89ms (Philips), or 4100/81.9ms (GE); a $90^o$ (Siemens), $78^o$ (Philips), or $77^o$ (GE) flip angle; mutliband factor = 3; 96 directions: 6 directions at b = 500, 15 directions at b = 2000, and 60 directions at b = 3000, resulting in $(1.7mm)^3$ isotropic resolution. Diffusion-weighted and functional scans were preceded by fieldmap scans to later correct for $B_0$ inhomogeneities.

#### Imaging data processing

For full details concerning the ABCD Study's MRI data processing pipeline, see Halger et al. (Hagler et al., 2019). Related code can be found on GitHub at https://github.com/ABCD-STUDY.

*Structural MRI*

Scanner manufacturer-specific transformations were used to correct gradient distortions in T1w images (Jovicich et al., 2006; Wald et al., 2001), followed by bias field correction via a sparse spatial smoothing approach that is based on the assumption that white matter has uniform intensity across the brain (Ashburner and Friston, 2000; Sled et al., 1998). T1w images were then registered to a 1.0 mm isotropic, in-house reference template via rigid body alignment (Friston et al., 1995), followed by resampling. FreeSurver v5.3 was used to, then, reconstruct the cortical surface and segment subcortical regions (Dale et al., 1999; Fischl, 2012). The cortical surface was (Fischl et al., 2002)reconstructed includes brain extraction, tissue type segmentation, the creation of a cortical mesh from the skull-stripped cortical gray matter, on which any topological errors are corrected and from which the optimal pial and white matter surfaces are created (Dale et al., 1999). Pial and white matter surfaces are then registered nonlinearly to a surface-based atlas by aligning gyral and sulcal patterns in a spherical space (Fischl et al., 1999). The usual FreeSurfer pipeline was modified to skip intensity scaling and inhomogeneity correction, as they had already been performed.

From these aligned surfaces, cortical gray matter regions were labeled per the Desikan & Killiany atlas (Desikan et al., 2006). Subcortical segmentation was also performed using FreeSurfer, in volumetric space (Fischl et al., 2002). White matter tracts were labeled using AtlasTrack, using a probabilistic fiber atlas that specifies specific long-range tracts with prior probabilities and orientation descriptions (Hagler et al., 2009). T1-weighted images are registered nonlinearly to the fiber atlas (Friston et al., 1995) and then diffusion orientations (from DTI data, see below) from each participant are compared to the atlas's orientations, per fiber. This information is used to update the tract priors for individual fiber tract regions of interest (ROIs), after excluding gray matter and cerebral spinal fluid (Fischl et al., 2002).

Gray-to-white matter contrast was computed using intensity values ± 0.2mm from the gray-white boundary (perpendicular at each point) with the following formula:

$$\frac{white - gray}{white + gray} \div 2$$

Values were averaged within each cortical ROI. Cortical thickness, area, and volume were computed using FreeSurfer and averaged within each cortical ROI (Chen et al., 2012; Fischl and Dale, 2000; Joyner et al., 2009; Rimol et al., 2010, 2010).

*Diffusion-weighted MRI*

First, diffusion data were eddy current corrected using a model to predict distortion patterns across all diffusion volumes based on the orientation and dispersion across gradients, while limiting displacement in the phase encoding direction (Andersson and Sotiropoulos, 2016; Barnett et al., 2014; Rohde et al., 2004; Zhuang et al., 2006). In the phase encoding direction, displacements were modeled by combining spatial information with gradient orientation and strength using 12 parameters across all volumes. Dark slices due to head motion were excluded using a robust tensor fit after identifying these slices using log-transformed images and censoring frames based on the root mean squared residual error per frame (Basser et al., 1994). These censored frames were replaced with an interpolated frame, using the tensor fit in successive iterations. Motion correction was performed using rigid-body registration to align each volume with the eddy current-corrected tensor fit, and the diffusion gradient matrix was updated to account for head motion (Hagler et al., 2009; Leemans and Jones, 2009). Magnetic field inhomogeneities were corrected using pairs of b = 0 images with opposite phase encoding, aligned via nonlinear registration, to estimate the displacement field. This estimated volume is used to correct each frame across successive diffusion-weighted volumes (Holland et al., 2010) and registered to the participant's T1-weighted image after pre-alignment using a within-modality atlas. Diffusion images are then resampled to align with a T1-weighted image that has been rigidly aligned to an atlas brain, to achieve a standard orientation for the resulting diffusion-weighted data that is consistent across participants. The diffusion gradient matrix was updated again, with respect to head motion.

Diffusion tensors were fit using linear estimation of log-transformed diffusion-weighted signals (Basser et al., 1994), separately for the inner shell and full shell. Inner shell models were fit excluding frames with b > 1000s/mm$^2$ (i.e., using 6 directions at b = 500s/mm$^2$ and 15, at b = 1000s/mm$^2$) and, thus, correspond to traditional acquisitions that only use a single b-value. Full shell models were fit using all directions and all gradient strengths. Diffusion tensor imaging (DTI) measures including fractional anisotropy and mean, longitudinal, and transverse diffusivity were computed from these data (Alexander et al., 2007), as were metrics using restriction spectrum imaging (RSI) (White et al., 2014). Briefly, RSI takes advantage of the multiple b-values to estimate "restricted" and "hindered" diffusion, per voxel, based on diffusion distance. These two volume fractions reflect intracellular (restricted) and extracellular (hindered) diffusion, with unique fiber orientation densities of which there can be multiple per voxel. From these fractions, we use restricted normalized isotropic and directional estimates (RNI, RND), which are unitless measures that range from 0 to 1. Thus, RND reflects diffusion that proceeds along an axis, i.e., oriented diffusion, and is similar to FA, but unaffected by crossing fibers. All RSI and DTI measures were calculated per white matter (WM) tract ROI, subcortical ROI, and cortical ROI. Weighting factors are included for subcortical averages to mitigate partial volume effects from adjacent cerebral spinal fluid (CSF), using Tukey's bisquare function on values of MD relative to the median in each subcortical ROI (Tukey, 1960).

*Functional MRI*

Preprocessing for functional MRI (fMRI) started with motion correction via 3dvolreg in AFNI (Cox, 1996), which aligns each volume to the first across a run and provides per-TR estimates of head motion that are later used in single-subject analyses. Magnetic field inhomogeneities are corrected using the same procedure as described above for diffusion data preprocessing. Then, images were distortion corrected (Jovicich et al., 2006) and each run is aligned to an example volume from the middle of the fMRI runs using rigid-body coregistration. Fieldmaps are registered to the participant's T1-weighted image, again, following pre-alignment with a modality-specific atlas.

Functional connectivity estimates were computed via seed-based correlation, adapted for use with cortical surface ROIs (Seibert and Brewer, 2011; Van Dijk et al., 2010). In addition to the previously described preprocessing, resting-state fMRI preprocessing includes the removal of initial TRs; time course normalization via mean-centering; and regression of quadratic trends, motion time series, motion censoring time series (i.e., volumes with framewise displacement > 0.3mm), and mean + first derivative time series from CSF and WM; and band-pass filtering (0.009-0.08Hz) (Hallquist et al., 2013; Power et al., 2014; Satterthwaite et al., 2012).

Average preprocessed time series were extracted from each cortical (in surface space) and subcortical ROI (in volume space). Further censoring was performed after time series were extracted from each ROI at FD > 0.2mm for network connectivity and BOLD variance estimates, removing high-motion volumes and time periods with fewer than five contiguous volumes (post-censoring). Then, outlier volumes were identified based on the standard deviation (SD) per ROI across TRs and time points with SD greater than three times the mean absolute deviation above or below the median SD were excluded. BOLD variance is calculated from each ROI, across the time series, and reflects the magnitude of low-frequency BOLD fluctuations. Network correlations were computed as the average Fisher-transformed correlation of each region comprising the network, per the Gordon parcellation (Gordon et al., 2016). Connectivity was calculated within each cortical network, between each pair of cortical networks, and between each cortical network and each subcortical ROI.

#### Included data per modality

Participants were excluded from these analyses if they did not have imaging data collected at the 2-year follow-up visit. From this subset, data were further excluded based on image quality. Structural (sMRI) and diffusion-weighted (dMRI) data were included if a participant's T1- and diffusion-weighted images, respectively, met the ABCD-recommended criteria for inclusion. For T1-weighted images, quality assessments were based on motion, intensity inhomogeneity, white matter and pial surface estimation by Freesurfer, and susceptibility artifacts and exclusion was recommended if an image exhibited severe artifact in any of those categories. For diffusion-weighted images, quality assessments were based on residual $B_0$ distortion after processing, coregistration to the participant's T1, image quality, and segmentation quality, with exclusion recommended if an image exhibited severe artifact in any of those categories. For resting-state scans, quality assessments were based on the number of frames remaining after high-motion (i.e., framewise displacement > 0.3 mm) frames were censored and periods with fewer than 5 contiguous frames were excluded, coregistration to T1, $B_0$ distortion maps, Freesurfer tissue type segmentation quality, and presence of runs with fewer than 100 usable timepoints (after censoring and exclusion). We additionally excluded any resting-state fMRI (rs-fMRI) data from participants with fewer than 750 usable (i.e., low-motion) frames, or 10 low-motion minutes, across all runs, based on estimations of scan lengths necessary for reliable resting-state functional connectivity estimates (Birn et al., 2013; Noble et al., 2017). The final sample characteristics for the current study are described in Table 1. Because each structural, diffusion-weighted, and resting-state functional data are included based on their image quality, there are different numbers of subjects included for analyses with different imaging modalities (Supplemental Table 2).

Brain measures from sMRI scans include cortical thickness, cortical area, cortical volume, subcortical volume, gray-to-white matter contrast, and white matter volume. Brain measures from dMRI scans include fractional anisotropy, mean diffusivity, longitudinal diffusivity, transverse diffusivity, and both isotropic and directional intracellular diffusion (in white and gray matter). Brain measures from rs-fMRI scans include BOLD temporal variance, between-network functional connectivity, within-network functional connectivity, and subcortical-network functional connectivity.

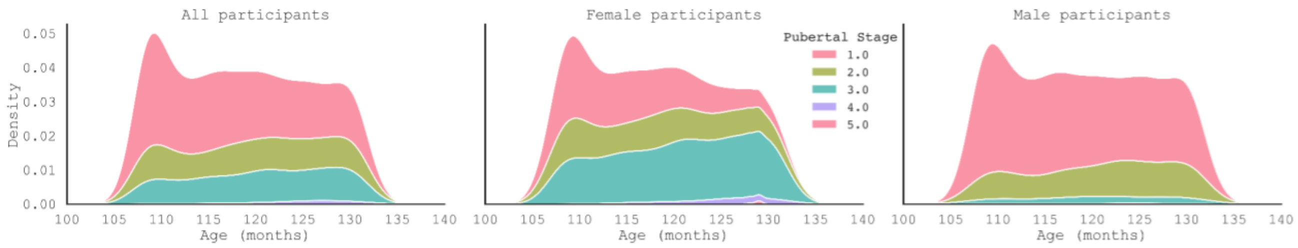


**Supplemental Figure 1.** Distributions of pubertal stage across participants and broken down by sex assigned at birth. 1 = prepubertal; 2 = early pubertal; 3 = mid-pubertal; 4 = late pubertal; 5 = post-pubertal

**Supplemental Table 2:** Sample Demographics Per Imaging Modality

| | All 2-Year Follow-Up | sMRI Pass QC | dMRI Pass QC | rs-fMRI Pass QC |
|---|---|---|---|---|
| Total N (%) | 7457 (100%) | 7115 (95%) | 6248 (84%) | 4119 (55%) |
| **Age at baseline (months)** | 118.74 ± 7.44 | 118.82 ± 7.45 | 118.90 ± 7.46 | 119.47 ± 7.55 |
| **Sex (F)** | 3437 | 3300 (46%) | 2922 (47%) | 2072 (50%) |
| ***Race & Ethnicity*** | | | | |
| Asian + Other | 918 | 870 | 732 | 481 |
| Hispanic | 1450 | 1383 | 1195 | 764 |
| Non-Hispanic Black | 981 | 924 | 787 | 416 |
| Non-Hispanic White | 4108 | 3938 | 3534 | 2458 |
| ***Household Income*** | | | | |
| >$100k | 1922 | 1817 | 1532 | 928 |
| $50 to $100k | 2082 | 1992 | 1756 | 1191 |
| <$50k | 2876 | 2760 | 2485 | 1715 |
| Don't Know / Refuse | 577 | 546 | 475 | 285 |
| ***Caregiver Education*** | | | | |
| Up to high school diploma, GED | 1147 | 1084 | 893 | 522 |
| Some college, associate's degree | 2233 | 2125 | 1866 | 1203 |
| Bachelor's degree | 2204 | 2125 | 1889 | 1308 |

**Supplemental Table 2:** Sample Demographics Per Imaging Modality

| | All 2-Year Follow-Up | sMRI Pass QC | dMRI Pass QC | rs-fMRI Pass QC |
|---|---|---|---|---|
| Graduate degree | 1863 | 1771 | 1591 | 1085 |
| Caregiver Marital Status | | | | |
| Married | 5192 | 4981 | 4436 | 3038 |
| Widowed | 60 | 56 | 48 | 34 |
| Divorced | 659 | 628 | 547 | 351 |
| Separated | 262 | 245 | 205 | 112 |
| Never Married | 850 | 797 | 668 | 378 |
| Refused to Answer | 43 | 41 | 35 | 15 |
| **MRI Scanner Manufacturer** | | | | |
| Siemens | 4539 | 4457 | 4116 | 2738 |
| GE Medical Systems | 2013 | 1810 | 1444 | 1040 |
| Philips Medical Systems | 905 | 848 | 688 | 341 |

*Note*. Differences in baseline demographic composition of the complete sample and the sample including individuals with high-quality MRI data collected at a 2-year follow-up appointment were assessed using Mann-Whitney U tests. Bold categories indicate significant differences at $\alpha < 0.05$; bold and italicized categories, at $\alpha < 0.01$. Within-category percentages that add to <100% are due to missing data and participants who declined to answer, i.e., in the case of household income. *MRI 2-Year Follow-Up* indicates the sample used in analyses presented here.

#### Associations between developmental variables

We assessed multicollinearity (via Variance Inflation Factor) and pairwise correlations (point biserial, Spearman where appropriate) between age, sex, and pubertal status across participants, to address any potential confounding of results. Age, sex, and puberty do exhibit multicollinearity with variance inflation factors of 4.53, 2.29, and 5.61, respectively. Likewise, pairwise correlations indicate that age and sex are significantly related (Point biserial $r = -0.036$, $p < 0.01$), age and puberty are significantly related (Spearman $r = 0.15$, $p < 0.01$), and puberty and sex are significantly related (Spearman $r = 0.42$, $p < 0.01$). Even though this relatedness is undoubtedly reflected in our results, neuroanatomical differences in patterns of heteroscedasticity between pubertal stage and each age and sex indicate that our results reflect distinct neurodevelopmental phenomena, too.

### Supplemental Results

#### Intra-individual variability

*Macrostructure changes*

Macrostructure changes are summarized in Table 3 and plotted in Figures 1 and 2A. The largest decreases of *cortical volume* were in posterior regions, both laterally and medially, including the rostral middle frontal gyrus, posterior cingulate gyrus, and right supramarginal gyrus. On the other hand, increases were seen in fewer cortical regions, including bilateral paracentral gyri, lateral orbitofrontal gyri, and lingual gyri. Conversely, *subcortical* structures all showed considerably larger increases in volume. *Cortical thickness* followed a similarly posterior trend, with the largest decreases in cuneus, precuneus, lingual gyri, and pericalcarine cortex, although with notable decreases in more rostral frontal areas such as the orbitofrontal gyrus. Increases in cortical thickness were few, only seen in bilateral entorhinal cortices. *Cortical area* showed lower magnitudes of change overall, including more anterior and medial increases (e.g., in cingulate, parahippocampal, and insular cortex) and posterior decreases (e.g., in parietal cortex, precuneus, and supramarginal gyrus). *White matter volume* increased across tracts, with the largest increases in the cingulum and parietal section of the superior longitudinal fasciculus; smallest, in the fornix and anterior thalamic radiations.

*Microstructure changes*

Microstructure changes are summarized in Table 3 and plotted in Figures 1 and 2B. Changes in microstructural were of the same magnitude as macrostructural changes and were largely heterogeneous depending on the measure and, to a lesser extent, the region. *Gray-to-white matter contrast* saw marked decreases in pre-, para-, and postcentral gyri, and increases only in more frontal regions (e.g., temporal pole, entorhinal cortex, rostral anterior cingulate cortex, frontal pole). *Fractional anisotropy* increased on the whole, with the greatest changes in the cingulum (both cingulate and parahippocampal portions), right inferior longitudinal fasciculus, and bilateral inferior fronto-occipital fasciculus. *Mean diffusivity* decreased overall, with the largest decreases in posterior portions of the superior longitudinal fasciculus, and the smallest decreases in the fornix and anterior thalamic radiations. *Longitudinal diffusivity* decreased most in the parietal portion of the superior longitudinal fasciculus, forceps minor, and in white matter between inferior and superior frontal cortex (frontal aslant tract). *Transverse diffusivity* decreased most in the forceps major, cingulum, and superior longitudinal fasciculus, and decreased least in the forceps minor and anterior thalamic radiation. I*sotropic restricted (i.e., intracellular) diffusion in white matter* increased most in the cingulum, decreased in the fornix. *Cortical isotropic intracellular diffusion* increased most in caudal middle frontal, superior frontal, pericalcarine, and transverse temporal gyri, but decreased in the banks of the superior temporal sulcus, pars orbitalis and triangularis of the inferior frontal gyrus, entorhinal cortex, and precentral gyrus. *Subcortical isotropic intracellular diffusion* only increased in the caudate, and decreased most in the nucleus accumbens, amygdala, and hippocampus. *Directional intracellular diffusion in white matter* increased most in the cingulum, internal capsule, and corona radiata, and least in the fornix. *Cortical directional intracellular diffusion* increased most in the right superior frontal and left caudal middle frontal gyri, but decreased most in the left superior frontal gyrus and the banks of the superior temporal sulcus. *Subcortical directional intracellular diffusion* only increased in the caudate and decreased most in bilateral amygdala, hippocampus, and accumbens.

*Functional changes*

Macrostructure changes are summarized in Table 3 and plotted in Figures 1 and 2C. Functional measures demonstrated a wider magnitude of change than did macro- and microstructural measures. *BOLD variance* increased most in left lateral occipital cortex, bilateral postcentral gyri, and bilateral entorhinal cortex, but decreased most in right lateral occipital cortex, pars orbitalis of the inferior frontal gyri, rostral middle frontal gyri, and rostral anterior cingulate cortex. *Within-network connectivity* increased most within the motor-hand and cingulo-parietal networks, but decreased most in the salience and ventral attention networks, as well as extra-network orbitofrontal and anterior temporal regions. *Between-network connectivity* increased most between the somatomotor hand network and somatomotor mouth, and frontoparietal networks, as well as between the salience network and both frontoparietal and cingulo-parietal networks. Between-network connectivity decreased the most between the ventral attention network and salience, default mode, and extra-network regions, as well as between the salience network and auditory, retrosplenial temporal, and dorsal attention networks. Finally, *connectivity between subcortical regions and cortical networks* (i.e., subcortical FC in Figure 1) increased the most between the nucleus accumbens and auditory, dorsal attention, and salience networks; between the brainstem and retrosplenial temporal and frontoparietal networks; between the right caudate and sensorimotor hand, retrosplenial temporal, and auditory networks; and between the left ventral diencephalon and auditory, retrosplenial, salience, and somatomotor hand networks. Subcortical-to-cortical network connectivity decreased most between the left putamen and cingulo-opercular, retrosplenial temporal, and frontoparietal networks; between the right putamen and visual and auditory networks; between the right thalamus and somatomotor mouth network; and between the left thalamus and both salience and cingulo-parietal networks.

#### Inter-individual variability

*Age*

Assessments of homogeneity of variance across four equal-sized bins in baseline age (9.0 -– 9.39; 9.4 -– 9.99; 10.0 -– 10.49; 10.5 -– 10.99) uncovered associations between age and between-individual *variability* of APΔ. Only gray matter volume, gray-to-white matter contrast, directional intracellular diffusion (in white matter), and BOLD variance displayed significant differences in inter-individual variability in brain development, and only in a few regions or tracts (<5%) with respect to baseline study age, compared to other brain measures. Pairwise, *post hoc* rank-sum test between binned ages revealed that, after correcting for multiple comparisons, no one age group consistently demonstrated significantly greater or lesser variability across all measures of APΔ or within the coarse groupings of macrostructure, microstructure, and function. Within individual measures, however, individuals ages 9.4 to 9.99 years at baseline demonstrated the most variability in gray-to-white matter contrast, WM directional intracellular diffusion, and BOLD variance, while monotonically increasing variability with increasing age was seen for gray matter volume (Supplemental Figure 2).

*Sex*

Sex differences in variance (i.e., heteroscedasticity with respect to sex) is displayed in Table 4, Figures 5-7, and Supplemental Figure 3). Cortical area displayed significantly different variability in 22% of regions between sexes; cortical volume, 13%; and white matter volume, 23%. Only one cortical region demonstrated sex differences in isotropic intracellular diffusion changes (left entorhinal cortex). Within-network functional connectivity displayed no significant sex-related heteroscedasticity, whereas changes in only one connection between cortical networks (Retrosplenial Temporal and Ventral Attention) and one connection between a subcortical region and a cortical network (Dorsal Attention, hippocampus) demonstrated sex differences in variability. Pairwise, *post hoc* rank-sum test between sexes revealed that, after correcting for multiple comparisons, neither male nor female participants demonstrated consistently greater variability across all measures of APΔ or within the coarse groupings of macrostructure, microstructure, and function. Rather, female participants demonstrated variability on some measures, while male participants demonstrated greater variability on others. Overall, gray matter volume, cortical area, and isotropic intracellular diffusion (GM) showed greater female-than-male variability across significantly heteroscedastic regions, while white matter volume and network connectivity (including subcortical connections) showed greater male-than-female variability across significantly heteroscedastic regions (Supplemental Figure 3).

*Puberty*

Puberty-related differences in variance (i.e., heteroscedasticity with respect to pubertal stage) is displayed in Table 4, Figures 5-7, Supplemental Table 3, and Supplemental Figures 4, 5, and 7. Across this sample, distributions of participants in “prepubertal”, “early puberty”, and “midpubertal” stages are approximately equal (albeit with different proportions of male and female participants within each stage, as seen in Supplemental Figure 1), with <100 participants considered “late pubertal”. Thus, heteroscedasticity was assessed between participants in “prepubertal”, “early puberty”, and “midpubertal” stages. Gray matter volume, cortical area, white matter tract volume, transverse diffusivity, isotropic intracellular diffusion (in gray and white matter) displayed the most significant heteroscedasticity with respect to pubertal status, in 11 to 40% of regions or tracts across the brain. On the other hand, white matter fractional anisotropy, mean diffusivity, and transverse diffusivity each displayed no significant heteroscedasticity, as did between-network connectivity. Pairwise, *post hoc* rank-sum tests in global trends per measure (i.e., including both significantly heteroscedastic regions and nonsignificant regions) revealed that mid-pubertal individuals display greater variance than early- or pre-pubertal individuals across functional measures ($p < 0.01$) but not across measures of macro- or microstructural change, while longitudinal diffusivity demonstrated significantly greater variability in pre-pubertal individuals than early- or mid-pubertal individuals.

Variance in APΔ across pubertal stages at ages 10-11 and 11-12 (i.e., the second and third waves of data collection) was not assessed, though some brain measures are heteroscedastic across those variables (see section 3.2.2.).

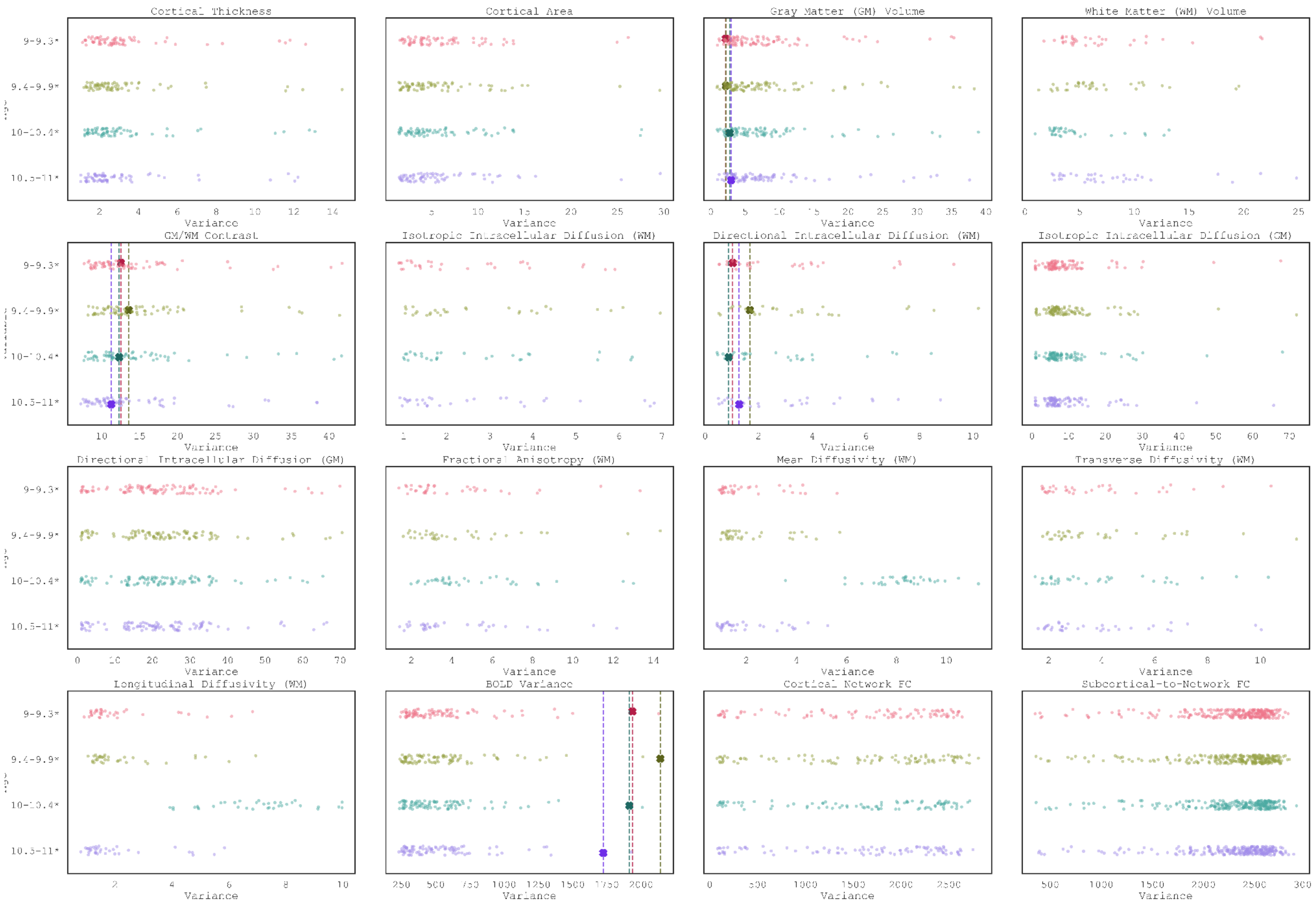


**Supplemental Figure 2.** Distributions of variance, per brain region, across age bins. Significantly heteroscedastic regions are marked by more opaque dots; not significantly heteroscedastic, by more transparent dots. The mean variance across significantly heteroscedastic brain regions, per measure, is denoted with a dashed line. Abbreviations: GM = gray matter; WM = white matter; FC = functional connectivity.

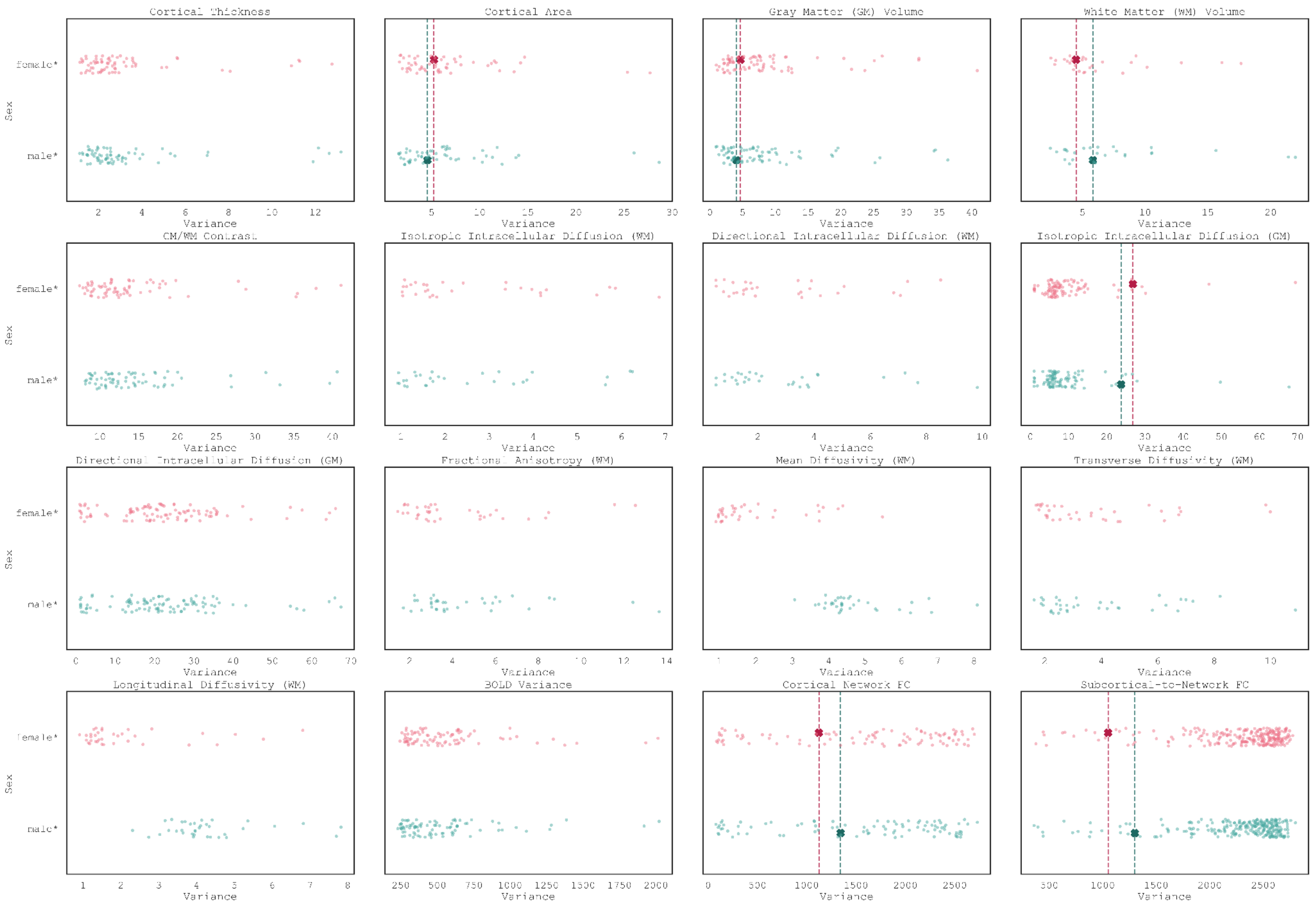


**Supplemental Figure 3.** Distributions of variance, per brain region, between sexes. Significantly heteroscedastic regions are marked by more opaque dots; not significantly heteroscedastic, by more transparent dots. The mean variance across significantly heteroscedastic brain regions, per measure, is denoted with a dashed line. Abbreviations: GM = gray matter; WM = white matter; FC = functional connectivity.

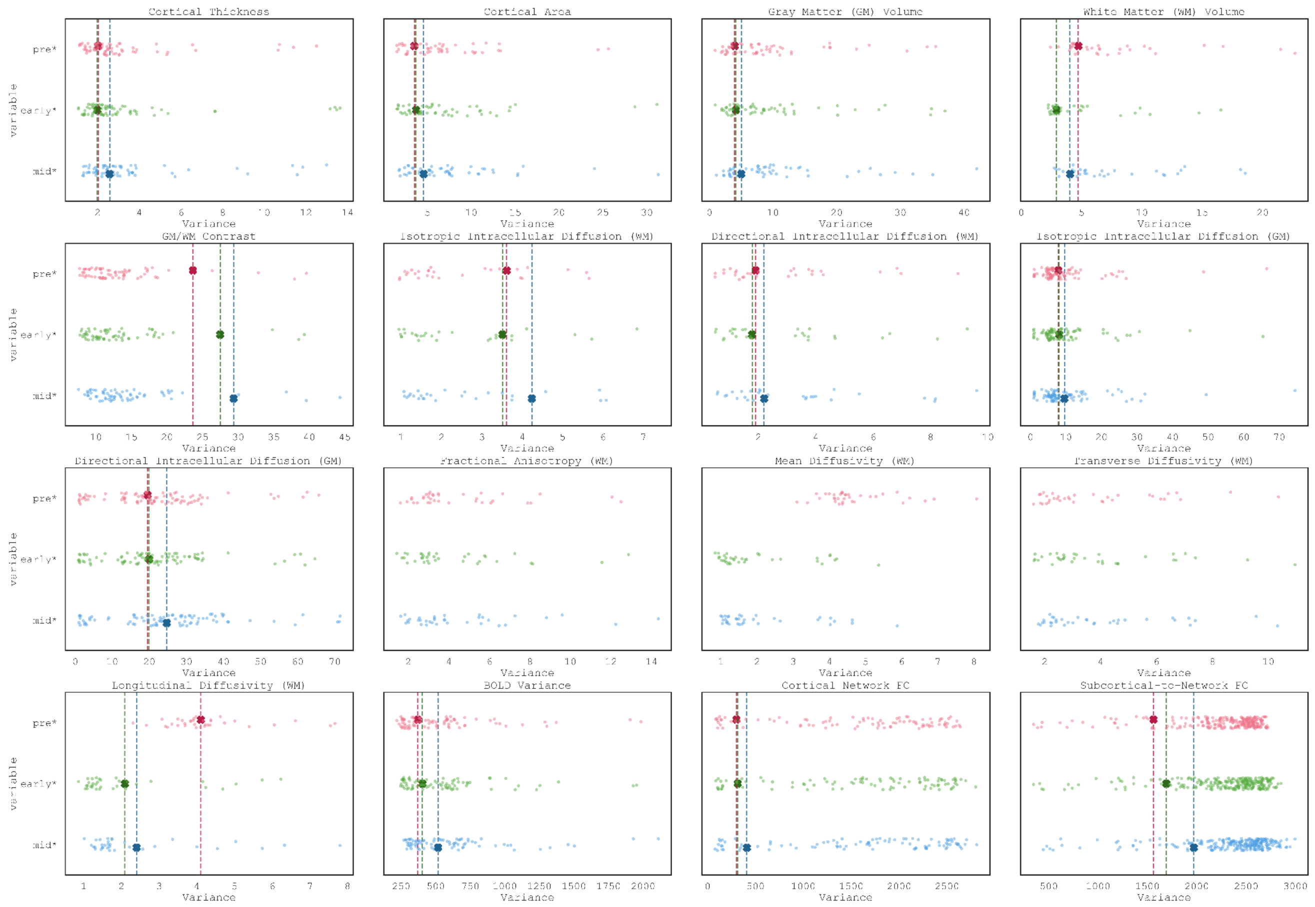


**Supplemental Figure 4.** Distributions of variance, per brain region, across pre-pubertal, early puberty, mid-pubertal, and late puberty participants. Significantly heteroscedastic regions are marked by more opaque dots; not significantly heteroscedastic, by more transparent dots. The mean variance across significantly heteroscedastic brain regions, per measure, is denoted with a dashed line. Late puberty is plotted, but as there are only 99 participants in the whole ABCD Study sample who were in "late puberty" at study enrollment these participants were not included in any analyses reported here and, thus, do not have mean lines plotted. Abbreviations: GM = gray matter; WM = white matter; FC = functional connectivity.

**Supplemental Table 3.** Mean variance across significantly heteroscedastic brain regions, per imaging measure, for participants in each pubertal stage at study enrollment.

| Measure | Prepubertal | Early puberty | Midpubertal |
|---|---|---|---|
| | $s^2$ | $s^2$ | $s^2$ |
| *Cortical Area* | 6.79 | 7.26 | 7.60 |
| *Cortical Thickness* | 3.13 | 3.28 | 3.48 |
| *Gray Matter Volume* | 9.53 | 10.03 | 10.60 |
| *White Matter Volume* | 4.73 | 2.91 | 4.04 |
| GM-WM Contrast | 13.49 | 13.53 | 14.71 |
| *Isotropic Intracellular Diffusion (GM)* | 9.48 | 9.52 | 10.46 |
| *Directional Intracellular Diffusion (GM)* | 21.68 | 21.53 | 25.13 |
| Fractional Anisotropy | 4.52 | 4.26 | 4.60 |
| Mean Diffusivity | 4.72 | 2.02 | 2.13 |
| *Longitudinal Diffusivity* | 4.26 | 2.11 | 2.42 |
| Transverse Diffusivity | 3.84 | 3.98 | 4.01 |
| *Isotropic Intracellular Diffusion (WM)* | 3.62 | 3.51 | 4.24 |
| *Directional Intracellular Diffusion (WM)* | 2.87 | 2.96 | 3.32 |
| *BOLD Variance* | 371.09 | 403.82 | **517.47** |
| *Within-Network FC* | 135.89 | 147.24 | 149.12 |
| *Between-Network FC* | 1784.64 | 1825.82 | 1858.46 |
| *Subcortical-Network FC* | 2261.50 | 2282.94 | 2356.42 |

*Note*: *There are <100 participants in this sample in "late puberty" at ages 9-10 years and, thus, they were not included in calculations of heteroscedasticity across pubertal stages. Bold indicates the stage with the greatest variability, based on *post hoc* rank-sum tests ($p < 0.01$). Measures with pubertal heteroscedasticity are indicated with an italicized label. Abbreviations: GM = gray matter; WM = white matter; FC = functional connectivity; BOLD = blood-oxygen-level-dependent.

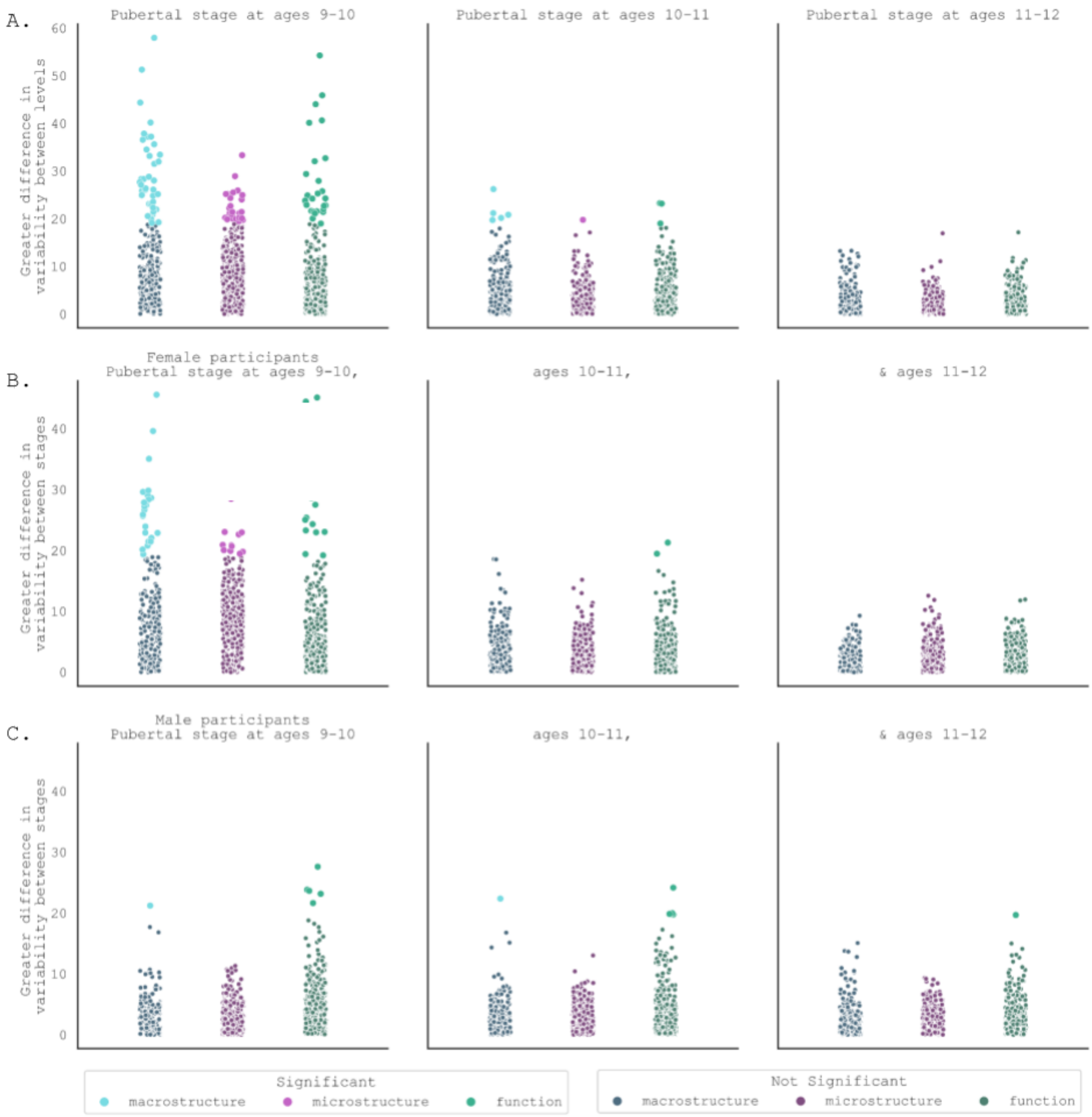


**Supplemental Figure 5.** Distributions of F-K statistics indicating heteroscedasticity in imaging measures with respect to pubertal stages. Significantly heteroscedastic regions/measures are brighter; not significantly heteroscedastic, darker. Top row: all participants. Middle row: female participants . Bottom row: male participants.

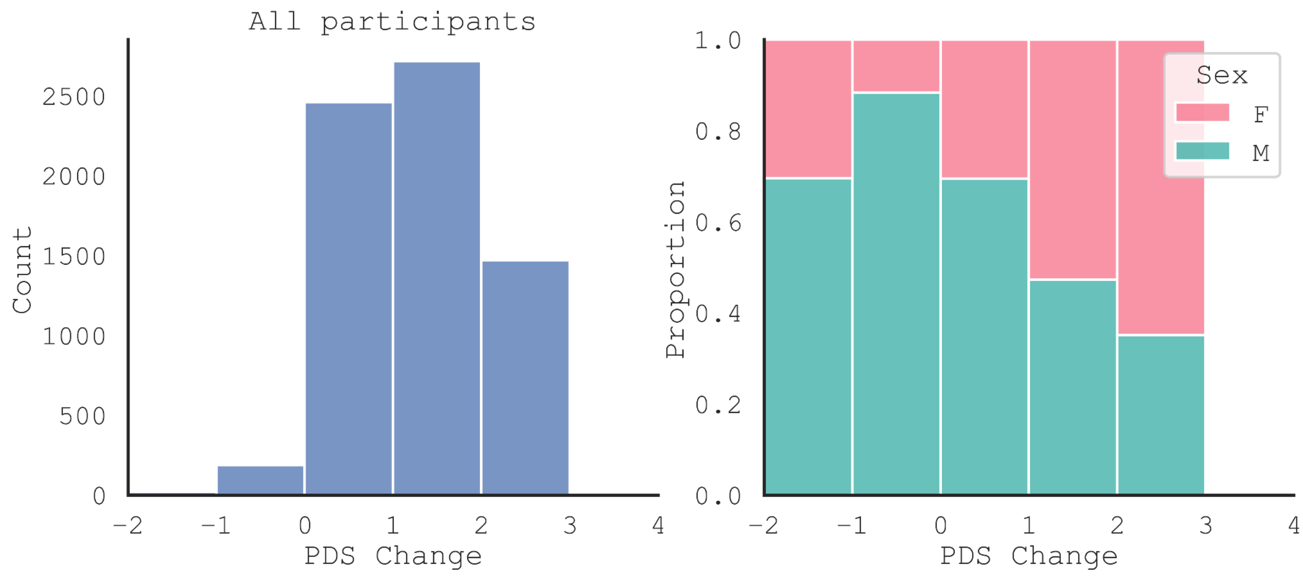


**Supplemental Figure 6.** Changes in PDS-determined pubertal stage across data collection time points for (left) all participants, (middle) female participants and (right) male participants. Note: annualized percent change presented in the manuscript was calculated between baseline and year 2 (as MRI data collection is every 2 years), corresponding to the blue bars on the plots.

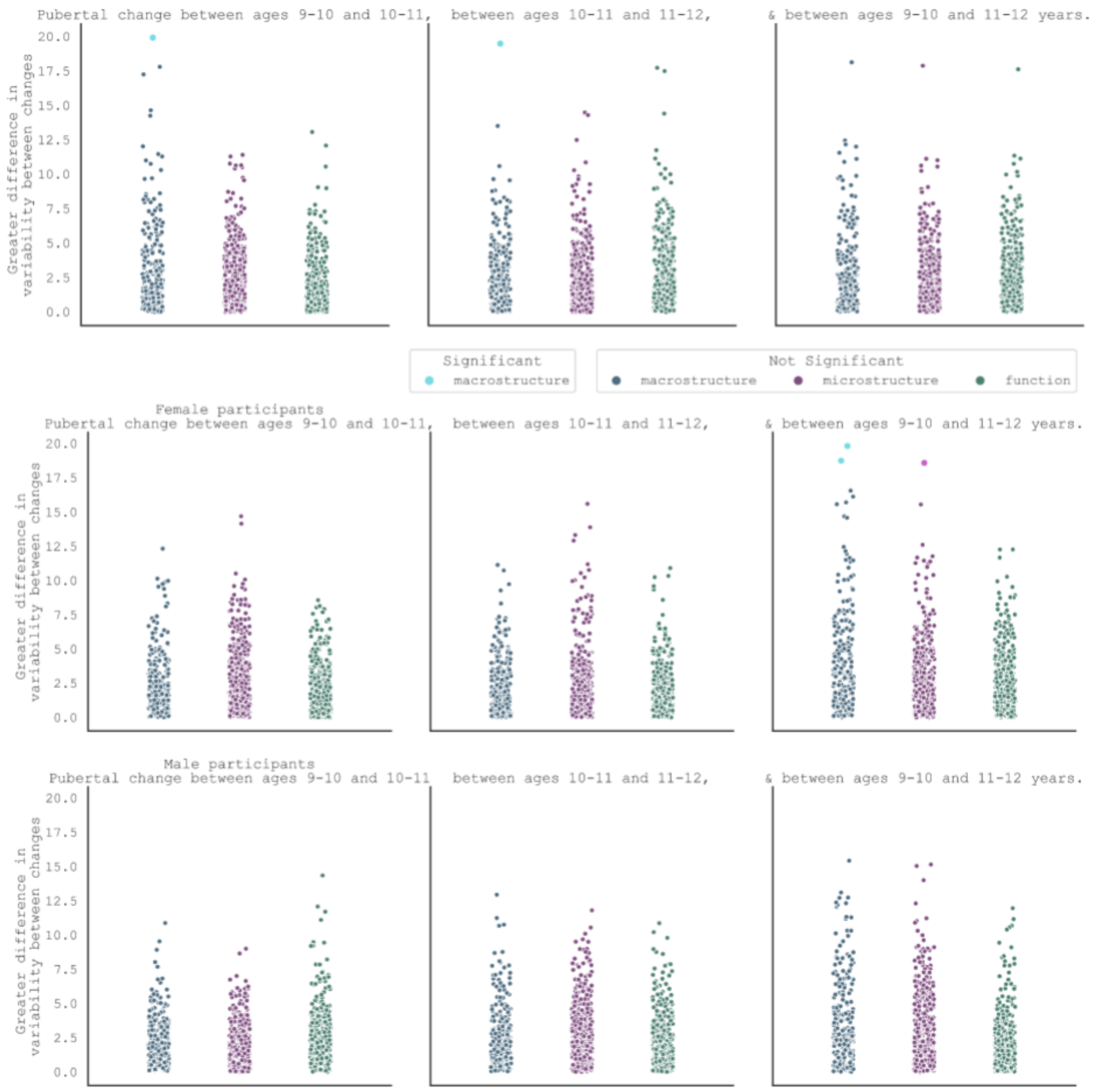


**Supplemental Figure 7.** Distributions of F-K statistics indicating heteroscedasticity in imaging measures with respect to changes in pubertal stages. There were no APΔs significantly heteroscedastic with respect to changes in PDS scores across time points, in any region/tract/network of any imaging measure. Top row: all participants. Middle row: female participants. Bottom row: male participants.

Feldstein Ewing, S.W., Miranda-Dominguez, O., Nagel, B.J., Perrone, A.J., Sturgeon, D.T., Goldstone, A., Pfefferbaum, A., Pohl, K.M., Prouty, D., Uban, K., Bookheimer, S.Y., Dapretto, M., Galvan, A., Bagot, K., Giedd, J., Infante, M.A., Jacobus, J., Patrick, K., Shilling, P.D., Desikan, R., Li, Y., Sugrue, L., Banich, M.T., Friedman, N., Hewitt, J.K., Hopfer, C., Sakai, J., Tanabe, J., Cottler, L.B., Nixon, S.J., Chang, L., Cloak, C., Ernst, T., Reeves, G., Kennedy, D.N., Heeringa, S., Peltier, S., Schulenberg, J., Sripada, C., Zucker, R.A., Iacono, W.G., Luciana, M., Calabro, F.J., Clark, D.B., Lewis, D.A., Luna, B., Schirda, C., Brima, T., Foxe, J.J., Freedman, E.G., Mruzek, D.W., Mason, M.J., Huber, R., McGlade, E., Prescot, A., Renshaw, P.F., Yurgelun-Todd, D.A., Allgaier, N.A., Dumas, J.A., Ivanova, M., Potter, A., Florsheim, P., Larson, C., Lisdahl, K., Charness, M.E., Fuemmeler, B., Hettema, J.M., Maes, H.H., Steinberg, J., Anokhin, A.P., Glaser, P., Heath, A.C., Madden, P.A., Baskin-Sommers, A., Constable, R.T., Grant, S.J., Dowling, G.J., Brown, S.A., Jernigan, T.L., Dale, A.M., 2019. Image processing and analysis methods for the Adolescent Brain Cognitive Development Study. NeuroImage 202, 116091. https://doi.org/10.1016/j.neuroimage.2019.116091

Hallquist, M.N., Hwang, K., Luna, B., 2013. The nuisance of nuisance regression: spectral misspecification in a common approach to resting-state fMRI preprocessing reintroduces noise and obscures functional connectivity. Neuroimage 82, 208–225.

Holland, D., Kuperman, J.M., Dale, A.M., 2010. Efficient correction of inhomogeneous static magnetic field-induced distortion in Echo Planar Imaging. Neuroimage 50, 175–183.

Jovicich, J., Czanner, S., Greve, D., Haley, E., van Der Kouwe, A., Gollub, R., Kennedy, D., Schmitt, F., Brown, G., MacFall, J., 2006. Reliability in multi-site structural MRI studies: effects of gradient non-linearity correction on phantom and human data. Neuroimage 30, 436–443.

Joyner, A.H., Bloss, C.S., Bakken, T.E., Rimol, L.M., Melle, I., Agartz, I., Djurovic, S., Topol, E.J., Schork, N.J., Andreassen, O.A., 2009. A common MECP2 haplotype associates with reduced cortical surface area in humans in two independent populations. Proc. Natl. Acad. Sci. 106, 15483–15488.

Leemans, A., Jones, D.K., 2009. The B‐matrix must be rotated when correcting for subject motion in DTI data. Magn. Reson. Med. Off. J. Int. Soc. Magn. Reson. Med. 61, 1336–1349.

Noble, S., Spann, M.N., Tokoglu, F., Shen, X., Constable, R.T., Scheinost, D., 2017. Influences on the Test–Retest Reliability of Functional Connectivity MRI and its Relationship with Behavioral Utility. Cereb. Cortex 27, 5415–5429. https://doi.org/10.1093/cercor/bhx230

Power, J.D., Mitra, A., Laumann, T.O., Snyder, A.Z., Schlaggar, B.L., Petersen, S.E., 2014. Methods to detect, characterize, and remove motion artifact in resting state fMRI. NeuroImage 84, 320–341. https://doi.org/10.1016/j.neuroimage.2013.08.048

Rimol, L.M., Panizzon, M.S., Fennema-Notestine, C., Eyler, L.T., Fischl, B., Franz, C.E., Hagler, D.J., Lyons, M.J., Neale, M.C., Pacheco, J., Perry, M.E., Schmitt, J.E., Grant, M.D., Seidman, L.J., Thermenos, H.W., Tsuang, M.T., Eisen, S.A., Kremen, W.S., Dale, A.M., 2010. Cortical thickness is influenced by regionally specific genetic factors. Biol. Psychiatry 67, 493–499. https://doi.org/10.1016/j.biopsych.2009.09.032

Rohde, G.K., Barnett, A.S., Basser, P.J., Marenco, S., Pierpaoli, C., 2004. Comprehensive approach for correction of motion and distortion in diffusion‐weighted MRI. Magn. Reson. Med. Off. J. Int. Soc. Magn. Reson. Med. 51, 103–114.

Satterthwaite, T.D., Wolf, D.H., Loughead, J., Ruparel, K., Elliott, M.A., Hakonarson, H., Gur, R.C., Gur, R.E., 2012. Impact of in-scanner head motion on multiple measures of functional connectivity: Relevance for studies of neurodevelopment in youth. NeuroImage 60, 623–632. https://doi.org/10.1016/j.neuroimage.2011.12.063

Seibert, T.M., Brewer, J.B., 2011. Default network correlations analyzed on native surfaces. J. Neurosci. Methods 198, 301–311.

Sled, J.G., Zijdenbos, A.P., Evans, A.C., 1998. A nonparametric method for automatic correction of intensity nonuniformity in MRI data. IEEE Trans. Med. Imaging 17, 87–97.

Tukey, J.W., 1960. A survey of sampling from contaminated distributions. Contrib. Probab. Stat. 448–485.

Van Dijk, K.R., Hedden, T., Venkataraman, A., Evans, K.C., Lazar, S.W., Buckner, R.L., 2010. Intrinsic functional connectivity as a tool for human connectomics: theory, properties, and optimization. J. Neurophysiol. 103, 297–321.

Wald, L., Schmitt, F., Dale, A., 2001. Systematic spatial distortion in MRI due to gradient non-linearities. NeuroImage 6, 50.

White, N.S., McDonald, C., Farid, N., Kuperman, J., Karow, D., Schenker-Ahmed, N.M., Bartsch, H., Rakow-Penner, R., Holland, D., Shabaik, A., Bjørnerud, A., Hope, T., Hattangadi-Gluth, J., Liss, M.,

Parsons, J.K., Chen, C.C., Raman, S., Margolis, D., Reiter, R.E., Marks, L., Kesari, S., Mundt, A.J., Kane, C.J., Carter, B.S., Bradley, W.G., Dale, A.M., 2014. Diffusion-weighted imaging in cancer: Physical foundations and applications of Restriction Spectrum Imaging. Cancer Res. 74, 4638–4652. https://doi.org/10.1158/0008-5472.CAN-13-3534

Zhuang, J., Hrabe, J., Kangarlu, A., Xu, D., Bansal, R., Branch, C.A., Peterson, B.S., 2006. Correction of eddy-current distortions in diffusion tensor images using the known directions and strengths of diffusion gradients. J. Magn. Reson. Imaging Off. J. Int. Soc. Magn. Reson. Med. 24, 1188–1193.
